# Urgency, Leakage, and the Relative Nature of Information Processing in Decision-making

**DOI:** 10.1101/706291

**Authors:** Jennifer S. Trueblood, Andrew Heathcote, Nathan J. Evans, William R. Holmes

## Abstract

Over the last decade, there has been a robust debate in decision neuroscience and psychology about what mechanism governs the time course of decision making. Historically, the most prominent hypothesis is that neural architectures accumulate information over time until some threshold is met, the so-called Evidence Accumulation hypothesis. However, most applications of this theory rely on simplifying assumptions, belying a number of potential complexities. Is changing stimulus information perceived and processed in an independent manner or is there a relative component? Does urgency play a role? What about evidence leakage? Although the latter questions have been the subject of recent investigations, most studies to date have been piecemeal in nature, addressing one aspect of the decision process or another. Here we develop a modeling framework, an extension of the Urgency Gating Model, in conjunction with a changing information experimental paradigm to simultaneously probe these aspects of the decision process. Using state-of-the-art Bayesian methods to perform parameter-based inference, we find that 1) information processing is relative with early information influencing the perception of late information, 2) time varying urgency and evidence accumulation are of roughly equal importance in the decision process, and 3) leakage is present with a time scale of ~200-250ms. To our knowledge, this is the first comprehensive study to utilize a changing information paradigm to jointly and quantitatively estimate the temporal dynamics of human decision-making.

---

Decades of research on the cognitive and neural processes involved in decision-making have converged on general agreement that information is sequentially sampled over time with a decision triggered when a threshold level is reached. However, how those samples are incorporated into the decision making process has been the source of much debate. Two seemingly distinct approaches have emerged. The conventional view is that evidence is accumulated or integrated over time until a threshold or decision bound is met. A second view, which has emerged more recently in the decision neuroscience literature, is that, rather than being integrated, incoming information is weighted by a temporally increasing “urgency signal” that magnifies the value of evidence as time progresses. Further complicating this debate is the intertwined question of whether, or to what extent, leakage and dependencies in information processing impact the decision process. In this article we develop a joint computational modeling and experimental framework to quantitatively assess the relative prominence of time varying urgency and integration in the decision process. Given that this question is intimately entangled with issues associated with evidence leakage and the potentially relative nature of information processing (which may be more important when evidence changes over time), we jointly consider all of these factors in our approach to ensure the absence of one factor does not lead to false conclusions about another.

The most heavily studied of these decision theories, the Evidence Accumulation Model (EAM), is supported by more than 50 years of behavioral research (Brown & Heathcote, 2008; Van Zandt, Colonius, & Proctor, 2000; Ratcliff & Smith, 2004; Dutilh et al., 2018; Stone, 1960) as well as neural recordings (Bollimunta, Totten, & Ditterich, 2012; Churchland et al., 2011; Cassey, Heathcote, & Brown, 2014). The more recently developed Urgency Gating Model (UGM), is similarly supported by both behavioral results (Cisek, Puskas, & El-Murr, 2009; Carland, Thura, & Cisek, 2015; Ditterich, 2006a; Drugowitsch, Moreno-Bote, Churchland, Shadlen, & Pouget, 2012) and neural recordings (Churchland, Kiani, & Shadlen, 2008; Thura & Cisek, 2016a). Furthermore, each view is grounded in optimality arguments. Simple random walk EAM’s are optimal in a stable environment in the sense of requiring the fewest evidence samples to achieve a given level of accuracy (Bogacz, Brown, Moehlis, Holmes, & Cohen, 2006). On the other hand, the UGM and related models that assume a temporally collapsing bound are suggested to optimize the time discounted reward rate, which accounts for not only the explicit rewards and penalties associated with a decision, but also the time costs associated with prolonged deliberation (Thura, Beauregard-Racine, Fradet, & Cisek, 2012; Drugowitsch et al., 2012).

The evidence for these different hypotheses is varied. Decades of behavioral observations and model comparisons (for reviews see Donkin & Brown, 2018; Ratcliff & McKoon, 2008) have shown EAM’s both qualitatively and quantitatively account for choice and response time (RT) data in a variety of speeded decision tasks. The urgency gating model, which has been subjected to far less testing due to its relative recency, has been supported by qualitative arguments about the persistence of information during a decision and how changes of information influence those decisions (Thura et al., 2012; Cisek et al., 2009; Thura & Cisek, 2016b; Carland, Thura, & Cisek, 2015; Zhang, Lee, Vandekerckhove, Maris, & Wagenmakers, 2014). More recently, studies have been carried out to assess the validity of these phenomenological hypotheses by fitting various EAM and UGM models to quantitative RT data (Hawkins, Wagenmakers, Ratcliff, & Brown, 2015; Hawkins, Forstmann, Wagenmakers, Ratcliff, & Brown, 2015; Evans, Hawkins, Boehm, Wagenmakers, & Brown, 2017). Results of these studies have generally found that EAMs better account for quantitative aspects of data than UGMs. However, there is also evidence for the presence of urgency in specific scenarios involving non-human primates whose decisions are motivated by a well defined learned reward schedule (Hawkins, Forstmann, et al., 2015), in humans when interrogation and free response paradigms are combined in the same task (Palestro, Weichart, Sederberg, & Turner, 2018), as well as in contexts where humans are given training and feedback comparable to what is common in non-human primate experiments (Evans & Hawkins, 2019).

Although these studies have varied in their methods and conclusions, their goal typically has been to determine which mechanism (integration or urgency) is most likely, based on a set of observations. However, these mechanisms are not mutually exclusive of each other. Rather, as suggested in Thura and Cisek (2016b) and Thura (2015), the canonical integration and urgency models can be conceptualized as extreme cases of a continuum of models whose behavior is determined by critical cognitive parameters encoding the strengths of leakage and urgency. The Diffusion Decision Model (DDM), for example, is associated with a fixed level of urgency that does not change over time and little, if any leakage (i.e., an infinite leakage time constant). The canonical UGM, on the other hand, is typically associated with a strongly time dependent urgency signal and fast leakage with a time constant 50 < *τ* < 200 ms (Ditterich, 2006b; Thura & Cisek, 2014). It is important to note that these hypotheses differ not only in their assumptions about the time varying nature of urgency but also in their assumptions about the persistence of evidence (leakage). Thus the relative importance of urgency versus integration cannot be studied in a vacuum; the potential presence of leakage must be jointly considered.

Significant efforts have been dedicated to assessing the strength of time dependent urgency (or caution) and what the time constant associated with evidence leakage is (Table 1). These efforts have largely taken one of two basic approaches, qualitative and quantitative. In one approach, researchers have sought to generate distinguishing qualitative predictions from competing hypotheses and design experiments to test those predictions. However, this approach has a number of pitfalls, most notably that it can be difficult to prove that the predictions are in fact distinguishing. An alternative approach is to quantitatively fit models encoding these parameters to data and either perform model comparisons or make inferences based on those parameters. This approach also has a number of pitfalls; the models must be identifiable and experimental paradigms must generate sufficiently rich data to constrain parameter estimates. Unfortunately, many of the models of time varying urgency (see Hawkins, Forstmann, et al., 2015; Ditterich, 2006a, for example) are not identifiable (Evans, Trueblood, & Holmes, 2019). Furthermore, most such studies have relied on experiments where evidence is fixed over time while it has been previously suggested (Cisek et al., 2009; and further supported here) that such experiments are not sufficient to distinguish between these hypotheses.

**Table 1.** Literature using modeling to investigate time varying urgency and collapse bounds in decisions. For each study we focus on 1) whether fixed or changing information experiments were utilized, 2) how time varying urgency was encoded into the modeling, 3) whether leakage was considered, and 4) whether quantitative fitting of models to data was utilized. In the “Urgency Type” column: Restricted UGM refers to the UGM with (k = ∞, see Equ. (3) below), unrestricted UGM refers to the UGM where k is unrestricted, NonLin Gain indicates an accumulator model with a nonlinear gain applied (similar to the UGM). In the Experimental Paradigm column, fixed and changing indicate whether stimulus evidence was held or fixed or allowed to change within trials and we also indicate whether response deadlines were used. In the Fit to Data Column, “Restricted” indicates that quantitative fitting was utilized but that important parameters (e.g. leakage, threshold, nondecision time) where fixed. Note that distilling entire studies into a single line is impossible, thus this is intended only to be a synopsis of these four characteristics of each study.

| Reference | Exp. Paradigm | Urgency Type | Leakage | Fit to Data |
| --- | --- | --- | --- | --- |
| Cisek et al., 2009 | Changing | Restricted UGM | Fixed (200ms) | No |
| Thura et al., 2012 | Changing | Restricted UGM | Fixed (100ms) | No |
| Thura & Cisek, 2014 | Changing | Unrestricted UGM | No | Restricted |
| Carland, Thura, & Cisek, 2015 | Changing | Restricted UGM | Fixed (250ms) | Restricted |
| Carland, Marcos, et al., 2015 | Changing | Restricted UGM | Fixed (167ms) | Restricted |
| Thura & Cisek, 2016a | Changing | Unrestricted UGM | No | Restricted |
| Winkel et al., 2014 | Changing | Restricted UGM | Infinitely Fast | Yes |
| Hawkins, Wagenmakers, et al., 2015 | Fixed | Restricted UGM | Fixed (100ms) | Restricted |
| Evans et al., 2017 | Changing | Restricted UGM | Estimated | Yes |
| Ditterich, 2006a | Fixed | NonLin Gain | No | Yes |
| Murphy et al., 2016 | Fixed, Deadline | NonLin Gain | No | Yes |
| Hawkins, Forstmann, et al., 2015 | Fixed | NonLin CB | No | Yes |
| Evans & Hawkins, 2019 | Fixed | NonLin CB | No | Yes |
| Evans, Hawkins, & Brown, 2019 | Fixed, Deadline | NonLin CB | No | Yes |
| Malhotra et al., 2017a | Fixed | NonLin CB | No | Yes |
| Zhang et al., 2014 | Fixed | NonLin CB | No | Yes |
| Voskuilen et al., 2016 | Fixed | NonLin CB | No | Yes |
| Drugowitsch et al., 2012 | Fixed | NonLin CB | No | Yes |
| Palestro et al., 2018 | Fixed, Deadline | NonLin CB | No | Yes |
| Voss et al., 2019 | Fixed | NonLin CB | No | Yes |
| Our Study | Changing | Unrestricted UGM | Estimated | Yes |

Although experiments where information changes during the course of a decision are necessary, they raise another confounding issue. Holmes, Trueblood, and Heathcote (2016) previously demonstrated that when information changes over time, early evidence alters how subsequent evidence is processed. In other words, information may be processed in a temporally relative, dependent manner rather than an independent manner. To our knowledge, Holmes et al. (2016) and Holmes and Trueblood (2017) are the only previous studies to account for this complication (this idea was also discussed in Ratcliff, 1980, but never tested with data). However, those studies did not consider the effects of leakage nor time varying urgency.

Our goal here is to formalize the space of evidence accumulation models differing in urgency and leakage characteristics into a quantitative framework that accounts for the interplay between these entangled factors. We will use this framework to determine the extent to which integration, time varying urgency, and evidence leakage play a role in decision-making. To our knowledge, this is the first such attempt to use parameter estimation in conjunction with changing evidence paradigms to jointly estimate, in a quantitative way, the importance of these factors.

The framework is based on a simple model of neural activation corresponding to a decision variable in speeded choice that includes both integration and urgency. Rather than being a new model, it is an extension of the existing UGM hypothesis that takes a broader view of the form and role of the proposed “urgency signal”. Specifically, we identify a single dimensionless parameter that quantifies the shape of this urgency signal and dictates the characteristics of the resulting decision. In extreme regimes of this parameter, the resulting model asymptotically takes on pure integration or pure urgency characteristics, while in intermediate regimes both are present. When combined with an additional Ornstein-Uhlenbeck like leakage parameter, which has sometimes been included in integration models (Busemeyer & Townsend, 1993; Smith, 1995; Usher & McClelland, 2001) and always in UGM models (Cisek et al., 2009; Thura et al., 2012), this framework gives rise to a rich two parameter family of models that encodes the relative importance of urgency vs. integration and short vs. long sensory memory. Crucially, this framework allows for the possibility that a mix of these features may be at work.

As anticipated, we found that fixed evidence paradigms lead to parameter indeterminacy in this model, and thus are unlikely to be effective at detecting urgency or leakage related effects. We therefore utilize a changing information experimental paradigm in combination with this framework to try to infer what decision strategy people are actually using. This changing information paradigm, where we vary the strength and duration of presented information during the decision process, allows us to both assess leakage and urgency characteristics and determine whether information is processed in a relative or veridical fashion. Across four different experiments, we demonstrate that participants consistently exhibit moderate levels of both leakage and urgency characteristics, indicating peoples’ decision strategies occupy a middle ground between the canonical leak-less integration and leaky urgency gating mechanisms. Our results further demonstrate that information processing is relative (rather than veridical) in nature and that the strength and duration of early evidence strongly affects how later evidence is processed.

The rest of this article is structured as follows. First, we provide a survey of previous relevant literature on this topic. Next we derive the mathematical formulation of this generalized UGM (gUGM) model and demonstrate that it effectively generates a two parameter family of models that differ in the strength of urgency and level of leakage. Given that one of the central theoretical arguments in favor of time varying urgency is that it leads to optimal behavior, we assess to what extent it is in fact optimal. We then undertake a parameter recovery study to demonstrate that this model is estimable if changing evidence is used. Finally we discuss the changing evidence paradigm and report empirical results as well as the quantitative fits of the gUGM.

## Revisiting past approaches

As illustrated in Table 1, numerous studies utilizing modeling have investigated the extent to which decision processes involve some form of time varying urgency or evidence leakage. Even though these previous studies have similar goals, they have taken significantly different approaches.

One key difference between these studies is the type of experimental paradigms they use to probe the decision process. Some have used paradigms where sensory evidence stays fixed during the course of individual decisions while others have utilized paradigms where sensory evidence changes. As an example consider the standard Random Dot Motion (RDM) paradigm. A fixed evidence paradigm would consist of trials where the direction of motion stays the same across the trial (e.g., the coherent set moves to the left) whereas a changing evidence paradigm involves trials where the direction of motion changes during the course of a single trial (e.g., motion switches from left to right). This difference is important as it has been previously suggested that changing information paradigms may be necessary for distinguishing the dynamic properties (e.g., integration versus urgency) of decision-making (Cisek et al., 2009). The rational behind this suggestion is simple: paradigms involving dynamic evidence are more effective for investigating dynamic properties of decisions. More recently, studies have also begun to use explicit deadlines based on the hypothesis that deadlines promote a sense of urgency (e.g. Palestro et al., 2018).

A second key difference between these studies is whether they incorporated evidence leakage, which determines the persistence of information. Although all studies surveyed here investigate whether or to what extent urgency plays a role in decisions, leakage is another important determinant of the dynamics of a decision process. Whereas urgency measures how the amount of evidence needed to induce a decision changes over time, leakage determines how sensory evidence is weighted over time. In particular, higher leakage rates correspond to faster attenuation of old sensory evidence. Prior studies (Radillo, Veliz-Cuba, Josić, & Kilpatrick, 2017; Kilpatrick, Holmes, Eissa, & Josic, 2018) have demonstrated that leakage is an integral part of optimal decision policies, particularly when sensory evidence is dynamic. Some studies have not considered the potential effects of leakage at all. Others have incorporated it, but fixed its rate at different values corresponding to timescales that vary from 100-250ms. One study (to our knowledge) has attempted to estimate rates of leakage from data (Evans et al., 2017), though they concluded that leakage was not meaningful and estimated the leakage timescale to be longer than trial times.

A third key difference between these studies is the extent to which quantitative fitting is used to either compare EAM and UGM theories or explicitly estimate the values of critical decision parameters. Some, particularly early studies such as Cisek et al. (2009) and Thura et al. (2012), utilized qualitative comparisons between models and data to assess whether evidence accumulation or time varying urgency best explain observations. A positive aspect of this approach is that it forces researchers to consider how the predictions of models might differ and design experiments around those predictions. Although this approach is sound in theory, a key requirement for it to be useful is that the qualitative predictions of models must be provably distinct, which is a high bar that can be difficult to meet (see Appendix for an example of how this approach can lead to misleading conclusions). Another challenge with this approach (as discussed in Evans et al., 2017) is that it can lend itself to cherry picking data, relying on analysis of only a subset of experimental conditions rather than treating an experimental data set as a whole.

An alternative approach has been to quantitatively fit models to data with one of two goals: either to compare the quality of model - data agreement in an attempt to elevate one theory over another, or estimate model parameters in order to make parameter-based inferences. The benefits of this approach are that it more concretely assesses the ability of a model to account for observations and is less prone to cherry picking data as models are typically jointly fit to entire data sets rather than only subsets. It does, however, have a number of pitfalls. In cases where the goal is to contrast models on be basis of quality of fit, common model comparison measures (e.g., information criteria such as AIC, BIC, or DIC) are often used to “pick a winner”. Unfortunately, these test statistics have numerous known issues, depending critically on the balance they strike between complexity penalties and quality of fit. Further, a single number summary fails to fully capture the full richness of the data or may be strongly influenced by small features of the data (Lee et al., 2019). In cases where parameter-based inference is used, it is also necessary to verify that the model’s parameters are identifiable on the basis of the type of data available. This is particularly true when non-linear functional forms of decision bounds or gain functions are utilized since, as shown in Evans, Trueblood, and Holmes (2019), these nonlinearities typically increase the parameter indeterminacy or “sloppiness” (Gutenkunst et al., 2007; Holmes, 2015) of models.

A fourth difference between these studies, though we would argue this is less fundamental in nature, is the type of modeling framework they used. The models in this field largely fall into two basic categories. 1) Models where sensory evidence is multiplied by time varying urgency signal or gain function producing a decision variable that builds up toward a fixed bound. 2) Diffusion like models where a decision variable builds up toward a time varying bound. While in this article we focus on the use of a generalized Urgency Gating Model framework, models involving collapsing bounds are in many ways comparable and thus we consider them in this survey of the literature. We also note that for purposes of this study, we differentiate between restricted and unrestricted UGM variants. The reason for this will become more clear in subsequent sections. In short, the UGM proposes that sensory evidence is multiplied by an “urgency” function that increases linearly with time. The “Restricted UGM” cases fixes the intercept of that linear urgency function at 0. Although this may seem like a minor restriction, as will be shown later, this has a huge impact on the model and restricts the scope of its dynamics in a fundamental way.

As Table 1 shows, no study that we are aware of has used a dynamically changing evidence paradigm in combination with quantitative model fitting to jointly determine the level of time varying urgency and evidence leakage in the decision process. Evans et al. (2017) has come the closest to date, but they used the Restricted UGM and focused on the difference between UGM and DDM caused by the increasing variance of moment-to-moment evidence produced by the ramping up of the urgency signal. Although this is not solely a result of the restriction placed on the UGM, that restriction will amplify the issue.

Combining all of these elements into a single study is critical to probe the dynamics of decisions. Paradigms where evidence dynamically changes naturally have the potential to yield more insight into the decision dynamics. Quantitative fitting and parameter estimation allows the data to tell us what best accounts for observations. In addition, the joint estimation of urgency and leakage ensures that we are not biasing results by fixing one model characteristic in order to estimate the other. In this article, we apply all of these elements simultaneously to several new experiments. Next, we develop a mathematical generalization of the UGM that includes distinct parameters representing urgency and leakage model characteristics. We then demonstrate that, in combination with a changing evidence paradigm, this model is recoverable and thus we can utilize parameter estimation. Finally, we develop a changing evidence paradigm over four distinct experiments and use state-of-the-art Bayesian methods to fit the gUGM and estimate leakage and urgency parameters.

### The Generalized Urgency Gating Model

In its most general form, the gUGM hypothesis states that the decision variable 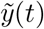 that tracks the temporal progress of a decision is influenced by two factors, a low pass filtered version of the integrated sensory evidence (*x*(*t*)) and a temporally increasing urgency signal (*U*(*t*)) (a restatement of Eqns. 1, 4 in Carland, Thura, & Cisek, 2015),

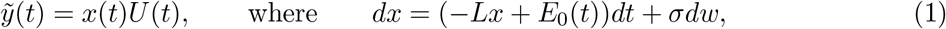

where 1/*L* encodes a leakage or low pass filter timescale and *E*(*t*) ~ *N*(*E*_0_(*t*), *σ*^2^) represents the time dependent and noisy evidence signal. The urgency signal takes the parametric form *U*(*t*) = *b* + *mt* where *b* and *m* represent, respectively, the baseline urgency and rate of increase of the urgency signal over time.

Thus far, in most cases simplifications of this model have been studied. Most studies of this model in human decision-making consider the restriction where *b* = 0, which as we will show greatly restricts the behavior of the model. In the primate neuroscience literature, this restriction has been relaxed but other assumptions have been made. In Thura, Cos, Trung, and Cisek (2014); Thura and Cisek (2016a, 2017) the full formulation of the urgency process was considered, but leakage was not as their stimuli do not possess perceptual noise. To our knowledge, the full formulation of this model with both the evidence leakage and the general urgency process (*b* ≠ 0) has not been quantitatively compared with data. This is critical since, as we will show, the fully general formulation of the UGM produces a parametric family of models that encode different types of decision strategies (e.g., DDM versus canonical UGM) in different regimes.

In order to clarify some of the salient properties of this model, we combine the expressions in Eqn. (1) using the product rule and non-dimensionalize the decision variable as follows (since in behavioral contexts, the values and units of this quantity are not observable). To begin, we calculate the differential of 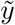

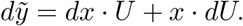

When dealing with stochastic processes one must take care in the type of product rule that is applied. However since *U* is not stochastic in this model, the simple product rule suffices and the Ito product rule is not necessary. An expression for *dx* is prescribed in Eqn. (1) and it is simple to calculate that *dU* = *mdt*. Substituting these into the above and simplifying yields the stochastic equation

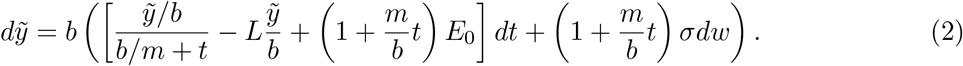

We now make the change of variables 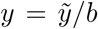 and define *k*:= *m/b*. From here on, we refer to this *k* as the “urgency ratio” since it describes the ratio of the background levels of urgency and the rate at which that urgency increases with time. Making these substitutions leads to a single stochastic differential equation describing the generalized model,

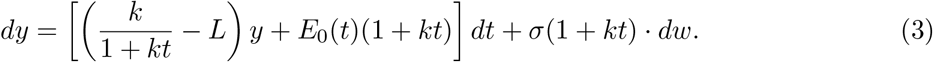

Here *y* represents the scaled (i.e., non-dimensional) decision variable and *k* = *m/b* is an “Urgency Ratio” parameter that will be the focus of much of the remaining discussion. In this formulation, there are two critical parameters that determine how this model behaves. *L* indicates the leakage rate, which is often described as setting the time constant of a low pass evidence filter, while *k* encodes the relative importance of time variation in the urgency signal. These two parameters selectively and orthogonally describe, respectively, the level of “memory” and importance of time varying urgency. In subsequent discussion and results, we expand on this statement and demonstrate that this framework provides a two-parameter continuum of models that unifies a diverse array of hypotheses previously proposed in decision neuroscience and psychology.

We do note briefly that a perceptual delay parameter is not included as was the case in Holmes et al. (2016). In that piecewise Linear Ballistic Accumulator (pLBA), a parameter *t_delay_* was incorporate to account for a potential temporal lag between the presentation of a new stimulus and the integration of that new information. While not exactly the same, the leakage in Equ. (1) produces a similar lag. That is, while *E*_0_, the stimulus strength, changes at the exact moment of the stimulus change, the resulting sensory evidence variable *x* will be subject to a lag due to leakage. We have thus chosen not to add an additional delay parameter.

In future discussion, we will be interested in the *k* = 0 and *k* → ∞ limits corresponding to “no urgency” (*m* = 0) and “standard urgency” (*b* = 0) respectively. The *k* = 0 limit is readily determined from Equ. (3) through substitution and yields the canonical Ornstein-Uhlenbeck process or drift diffusion model with leakage

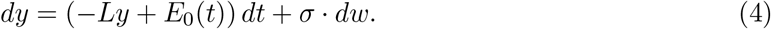

When *L* = 0, this leads to a standard diffusion process (Ratcliff, 1978) and thus we term the *k* = 0 limit the integration limit. Simple substitution cannot be used to determine the dynamics in the *k* = ∞ limit for mathematical reasons. However a similar derivation can be carried out in the case where the intercept of the urgency signal is *b* = 0, which is the standard UGM that has been studied in most cases. In this case, Equ. (2) becomes

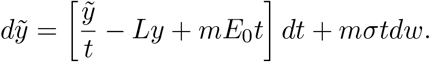

Recall that the moment to moment evidence is *E* ~ *N*(*E*_0_, *σ*^2^). Scaling this distribution by the constant *m* yields *mE* ~ *N*(*mE*_0_, (*mσ*)^2^). Thus the terms *mE*_0_ and *mσ* are essentially rescalings of the evidence and thus we simply fold *m* into the parameters *E*_0_ and *σ*. Dropping the ^~^ yields

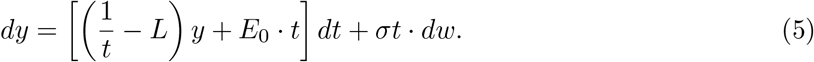

## The gUGM as a two parameter family of models

Two critical parameters in this gUGM determine the qualitative characteristics of its dynamics, the urgency ratio *k*, and the leakage rate *L*. The role of *L* is clear, it determines the level of “memory” the resulting stochastic process encodes. The other critical element of the UGM hypothesis is the presence of a temporally increasing urgency signal. The form of that urgency signal, which is encoded in the non-dimensional Urgency Ratio (*k*), modulates the behavior of the underlying model as well as the qualitative interpretation of the cognitive strategy it encodes. Consider two limiting cases, very small (*k* ~ 0) and very large (*k* ~ ∞). In the former case, a simple substitution of *k* = 0 leads to a reduced model (Equ. (4)) which is precisely an evidence integrator, where the value of *L* determines the presence / importance of leakage. In the other extreme, the resulting reduced model takes the form (Equ. (5)) of the “pure UGM”, which has been implemented and tested in Cisek et al. (2009); Thura et al. (2012).

More generally, the parameters (*k, L*) generate a two parameter family of models that differ in the level of “memory” (i.e., the inverse of leakage) and relative importance of urgency / integration. As illustrated in Figure 1a, in one quadrant of this modeling space recovers the canonical diffusion models and another the canonical pure UGM. Thus, while the urgency ratio encodes the relative importance of urgency gating versus evidence accumulation (*k* < 1 represents an integration dominated system while *k* > 1 is urgency dominated), it is the joint action of these two model components that determine the resulting decision strategy. Although this point has been alluded to in prior studies (Thura & Cisek, 2016b for example), the gUGM formalizes this model space and provides a path forward to jointly estimate these critical parameters from data.

**Figure 1.**
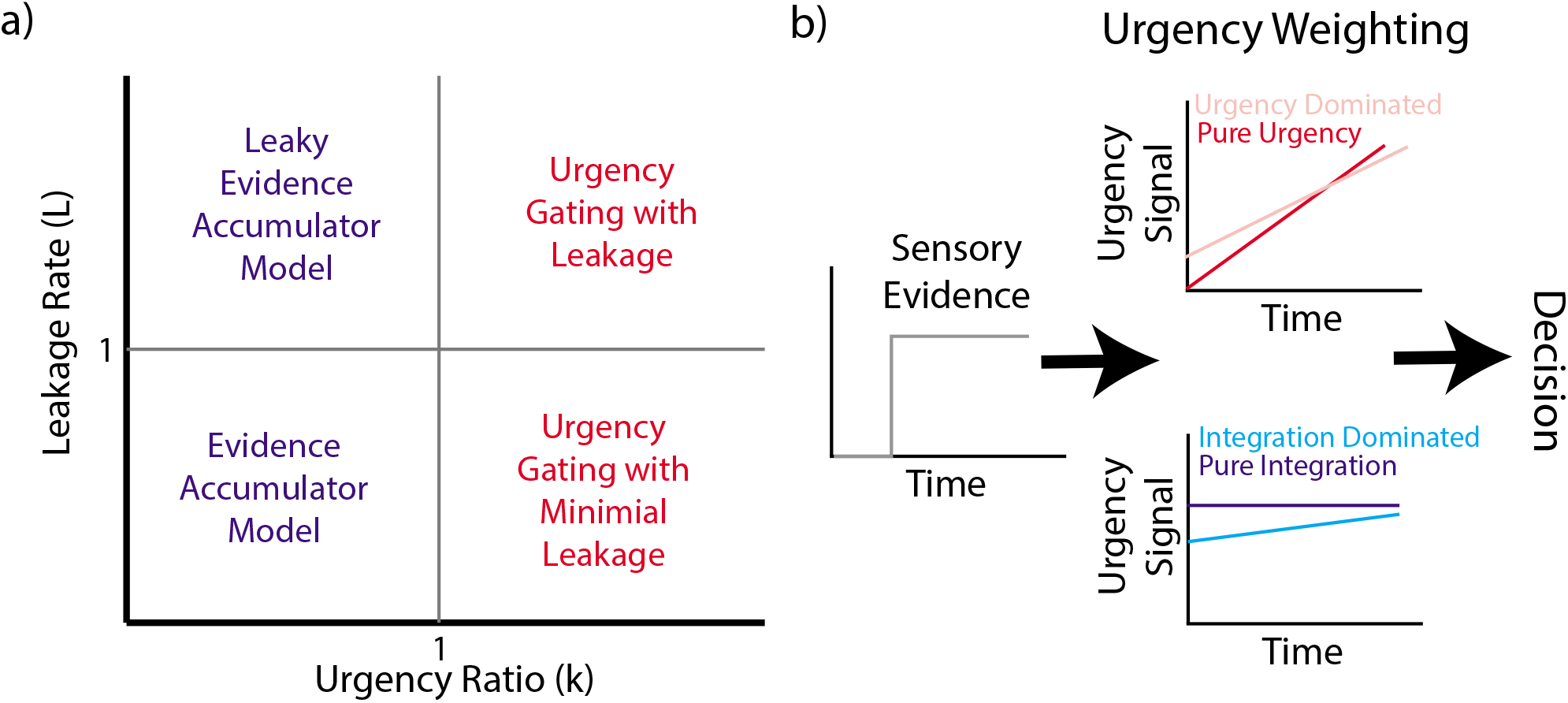
Synopsis of reduced model variants. **Panel a)** Schematic depiction of the two parameter family of models in the generalized UGM determined by the urgency ratio (k) and leakage rate (L). **Panel b)** Schematic description of the generalized UGM and the association between urgency signal shape and interpretation. Dark red (pure urgency) is associated with *k* = ∞, light red (urgency dominated) with *k* > 1, dark blue (pure integration) with *k* = 0, and light blue (integration dominated) with *k* < 1.

To demonstrate the effect of these two parameters on the shape of the decision variable over time, Figure 2 illustrates the simulated (and normalized) traces of the generalized model for various values of (*k, L*) with changing evidence (*E*_0_ = 1 for *t* < 1 and *E*_0_ = –1 for *t* > 1), where the panels highlighted in blue and green indicate canonical EAM (with very low *k*) and time varying urgency dominated (very high *k* with leakage corresponding to an effective timescale of 250ms) models. In every panel the straight grey lines with unit slopes (positive before the change and negative after) are plotted so that the rate of increase/decrease in the decision variable (plotted in black) can be compared to that of the DDM (i.e., a linear integrator with no leakage). In the top left panel it is evident that for the smallest value of *k* and no leakage the decision variable is indistinguishable from the DDM. In the pure UGM case with moderate leakage (red box, which corresponds to the canonical UGM studied to date), the initial phase of accumulation appears to be indistinguishable from a DDM. This, however, is not solely an effect of the urgency itself since pure urgency with longer leakage timescales yield different behavior. Rather, the interaction between urgency and leakage is responsible for this initial phase of linear integration. While the case highlighted in red may resemble an integrator prior to the change, the decision variable deviates dramatically from that of the DDM after the change. Thus the temporal characteristics of the decision variable of the gUGM varies considerably in different scenarios, particularly when changes of evidence are introduced. This observation also provides the first hint as to why changing information paradigms are likely more suitable to interrogate how integration versus urgency dominated the decision process may differ.

**Figure 2.**
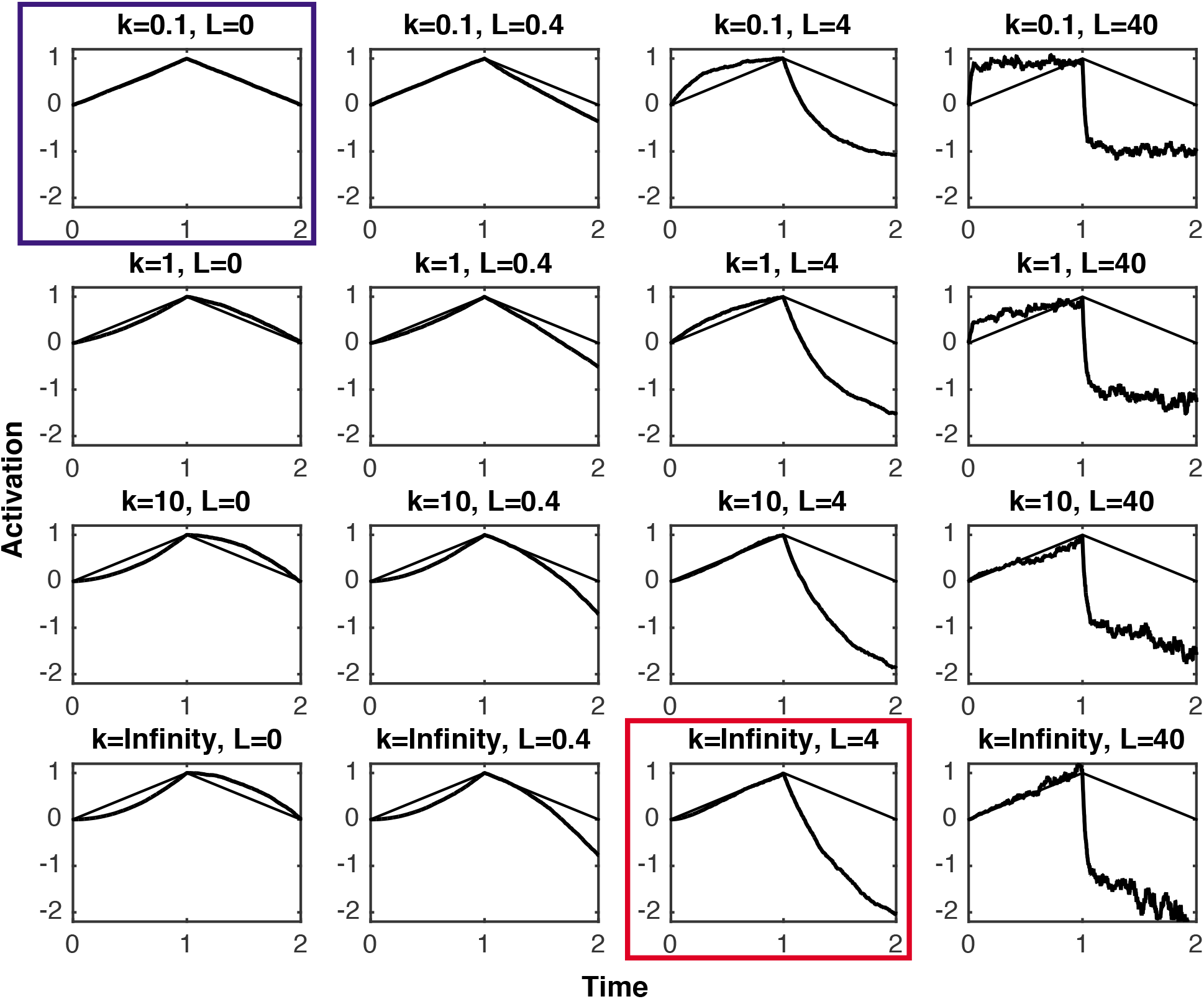
Simulated decision variable traces with changing information. Averaged simulation results for the generalized UGM with evidence that changes from *E* = 1 to *E* = −1 at *T* = 1. For each set of parameters, Eqn. (3) was simulated 100,000 times with a 1ms time step and the average trace is plotted. All simulations are run to a final time of *T* = 2 and simulations are normalized so that the mean end value is *ȳ*(*T* = 1) = 1 for each of side by side comparison. On each plot, grey lines with slope = 1 on *T* = [0,1] and slope = −1 on *T* = [1,2] are plotted so that the rate of increase in the decision variable can be compared to that of a linear integrator (i.e., the DDM). For reference, the panels encircled in blue / green correspond to pure integrator and pure UGM (with 250ms leakage constant), respectively. Recall that *τ* = 1/*L* is the effective timescale of evidence filtering / leakage.

### Investigating optimality with the gUGM

One of the central intuitive arguments in favor of some form of urgency signal (Thura et al., 2012) or time varying caution (Manohar et al., 2015; Drugowitsch et al., 2012) has been the supposition that it optimizes the decision makers time discounted reward rate, which accounts for both the reward associated with a decision as well as the time cost of making it. Thus, we next utilize this generalized model to assess whether there is an optimal level of urgency.

The reward-rate function

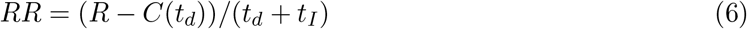

is often used to quantify the performance of a participant on a reward motivated experiment. Here, *R* represents the explicit reward (±1 for correct and incorrect responses) received on an individual trial, *t_d_* the decision time, and *t_I_* the waiting time / inter-trial time between successive trials. *C*(*t_d_*) represents the total cost of accumulating evidence during deliberation. This introduces two distinct costs associated with long deliberation times: 1) the time discounting through the term 1/(*t_d_* + *t_I_*) which reduces RR, and 2) the cost function *C*(*t_d_*) (investigated in Drugowitsch et al., 2012). The intuitive case for urgency modulation is, in part, that it reduces the cost associated with long deliberations (by preventing them) so that it maximizes accuracy subject to time cost.

Within this reward-rate framework, there are multiple ways to account for the explicit cost of accumulating evidence *C*(*t*). A general conceptualisation is in terms of a cost-per-unit-time function (*c*(*t*)), which quantifies the cost of accumulating evidence in a small unit of time *t* + Δ*t* rather than the total deliberation time. The two are related through *C′*(*t*) = *c*(*t*). We consider two extreme cases for the cost of accumulation. First, *C*(*t*) = 0, which corresponds to no cost of accumulation. In this case, the only effect of time on reward rate is time discounting due to deliberation time. In the second case we assume *c*(*t*) = *mt*, in which case *C*(*t*) = *m*/2 *t*^2^. We use this extreme case in which the cost per unit time linearly increases to determine the effect of this additional factor when compared to the former case. In Drugowitsch et al. (2012) it was suggested that this function *c*(*t*) likely initially increases somewhat linearly and subsequently saturates at larger times. However we chose to consider the two extreme ends of the spectrum here since numerous choices could in principle be made for this function. We chose a value of *m* = 1 for the cost slope. We did, however, vary this parameter as well, in results not shown, which lead to the same qualitative conclusions as discussed below.

Here we use this generalized framework to investigate the link between urgency and reward rate. For this study, we will consider level of caution (response threshold) and strength of urgency (urgency ratio) as cognitive parameters that could be potentially modulated or tuned through trial and error learning. To investigate this link, we design an *in silico* experiment mimicking a real experiment and assess the reward rate of *in silico* participants described by the model with different parameters. Since it has been suggested that the importance of urgency is most salient in cases where information changes during a decision or the difficulty of a decision varies unpredictably from trial to trial (Thura et al., 2012; Cisek et al., 2009; Thura & Cisek, 2016b; Carland, Thura, & Cisek, 2015), we designed the simulation experiment to consist of a range of difficulties with both fixed and changing information trials. For simplicity, we fix the value of leakage, the inter-trial interval length, and the specific form of the reward-rate function, and simulate the results of this experiment for a large collection of *in silico* participants that differ in their urgency and threshold parameters.

The *in silico* experiment we consider here consists of two types of trials, fixed information and changing information. Information in this context refers to the mean evidence strength (*E*_0_(*t*)) of a stimulus. In the fixed case, we assume *E*_0_ ~ *U*(0, 2) is a fixed value independent of time, that varies from trial to trial according to a uniform distribution. This distribution ensures that trials with a range of difficulties (*E*_0_ = 2 corresponds to very easy or strong sensory evidence for example, and *E*_0_ close to 0 corresponds to very hard) are considered. The second trial type is changing information. In this trial type, *E*_0_ takes on one fixed value prior to the change of information (which occurs at *t_s_* = 0.2 sec), and a separate value after, both of which are drawn from the same distribution as the fixed trials. Our *in silico* experiment consists of 10,000 copies of each of the two trial types to reduce the effects of randomness and to get a accurate representation of the aggregate reward rate. Real participants would of course not have this benefit, likely introducing significant noise into the RR landscape. Different values of the switch time and the upper limit on the distribution of difficulties were considered, all yielding similar qualitative results (not shown).

In order to map the influence of urgency and threshold on performance (as measured by reward), we generated *in silico* participants with a grid of values of each parameter. The threshold and urgency ratio were logarithmically sampled on the range (0.1, 10) and (0.1, 20) respectively with 50 values of each parameter. Logarithmic sampling was performed to ensure an equal distribution of small, medium, and large values of each parameter. In total, this leads to 2,500 *in silico* participants. The performance / reward rate of each of these *in silico* participants was assessed with results shown in Figures 6(a-d). Values of the remaining parameters (leakage, switch time on changing trials, inter-trial interval) as well as the form of the cost function were also varied, though only a representative sampling of results are shown since all simulations support the same conclusion, that there is no well defined optimal set of parameters.

Figures 6 (a-d) shows the relationship between the cognitive parameters (namely, response threshold and urgency) and reward rates in four different scenarios. The first, a base case with nominal parameter values, demonstrates that there is no well defined “optimal” parameter set that maximizes reward rate. Rather, there is a distributed band of parameter combinations that achieve comparable reward-rate values with a non-distinct peak at intermediate urgency-ratio values. Said another way, for every urgency strength there is a threshold value that achieves a nearly optimal reward rate. The remaining scenarios show that modulating the leakage rate, the inter-trial interval, and introducing a non-zero cost function *C*(*t_d_*) all lead to similar results. Although increased urgency does not appear to improve reward rate, it does tend to increase the range of thresholds that yield a reasonable reward rate (defined as greater than 50% of the maximal reward rate), thus making the model less sensitive to threshold levels. This is particularly evident when there is a short inter-trial interval, in which case the total time to achieve a reward becomes dominated by decision time rather than inter-trial time.

Alternative *in silico* experimental designs were also considered. In one alternative, rather than *E*_0_ taking on a distribution of values, we considered a design where it took on only two values corresponding to weak (*E*_0_ = 0.2) and strong (*E*_0_ = 2) evidence, similar to Malhotra, Leslie, Ludwig, and Bogacz (2017b). In a second alternative, we fixed the total experimental time (instead of fixing the number of trials performed). In this case, intermixed trials of different difficulty were performed until the time limit was reached (similar to Malhotra et al., 2017a), which introduces a global time penalty. In this case, rather than assessing the *reward rate* as a function of parameters, we calculated *total reward* over the experiment to assess the effect of parameters. Both experimental designs yielded similar qualitative results, that is, no well defined optimal parameters (results not shown).

Our results suggest that increasing urgency strength does not necessarily improve reward rate, even when information changes over time and difficulty is modulated between trials. However, we recognize that we investigated but one instantiation of the urgency gating hypotheses. For example, we instituted a linearly growing cost function (*C*(*t*)) while it has been proposed this function may saturate (Drugowitsch et al., 2012). That said, our results are similar to those of Malhotra et al. (2017a), where it was shown that collapsing-threshold models have a similarly broad band of “nearly optimal” parameters and that real participants seem to occupy a diverse range of locations in that band rather than all coveraging onto a single optimal state.

## Urgency, leakage and the need for changing information paradigms

This full generalization of the UGM describes a two parameter family of models whose properties are dictated by the urgency ratio *k* and leakage *L*. In the appropriate limits it gives rise to pure EAM models, pure UGM models, and models where each is differentially present. We next ask whether, and in what experimental design, the gUGM has the potential to serve as a diagnostic tool to assess the relative importance of these proposed cognitive mechanisms. That is, with an appropriate data set, could these critical parameters actually be jointly estimated? To address this, we performed a large scale parameter recovery experiment utilizing two types of stimulus information, fixed and changing.

To obtain a comprehensive assessment of the identifiability of the central parameters of Eqn. (3), we performed a large simulation study. We focused on four parameters (*k, L, v, a*), where *v* is the parameter representing the strength of evidence (e.g., *v* = *E*_0_). A large number of parameter sets (3000 in total) were randomly chosen using a latin hypercube design and for each resulting parameter set, a synthetic data set was simulated. For purposes of this study, we fixed the encoding and response time at *t_er_* = 0.3 and the trial to trial variability in *v* at *s* = 0.1. The four key parameters were varied across the following ranges:

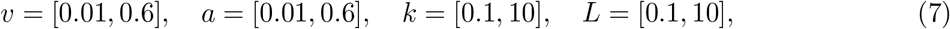

where “v” and “a” were sampled uniformly while “k” and “L” were sampled logarithmicaly to ensure equal coverage of both low and high values.

For each parameter set, two different data sets were generated, one with fixed evidence and the other with evidence that changes once during the trial. For the fixed evidence case, 1000 independent simulations were performed to generate a synthetic RT distribution. In this case, the evidence signal takes the value *E*_0_ = *v* for the full duration of the trial. For the switching evidence case, we simulated two synthetic trial types (T1 and T2). In T1, evidence remained unchanged for the entirety of the trial (*E*_0_ = *v* for the full duration). For T2, the stimulus changes at 0.2s after initiation (e.g. *E*_0_ = *v* for *t* G [0,0.2] and *E*_0_ = –*v* for *t* ∈ [0.2, end]).

We censored this collection of parameter sets to ensure only reasonable RT distributions were considered for further analysis. Non-switching RT data sets (all data for the fixed evidence case but only T1 data for the switching evidence case) had to meet the following criteria in order to be considered a “typical” response time distribution: the mean and median response times had to be between 0.4 and 2.5 seconds, the interquartile range was required to be between 0.1 and 2 seconds, the minimum RT was required to be below 1.5s, and the maximum was required to be greater than 0.5s. After applying these conditions, 926 and 1058 parameter sets (and their associated simulated RTs) remained in the constant evidence and changing evidence data sets respectively for further analysis.

Next, the generalized UGM was fit to each of these synthetic data sets to determine its ability to recover parameters. All recoveries were performed using Bayesian parameter estimation using Differential Evolution Markov chain Monte Carlo (DE-MCMC; Turner, Sederberg, Brown, & Steyvers, 2013) with 3N chains (where *N* is the number of free parameters in the model). In each case, a 1000 iteration burnin was used after which 2000 samples were obtained from the posterior. In the switching-evidence cases, the switch times were assumed to be known inputs to the model. Since this model does not have an analytical solution, we utilized the Probability Density Approximation (PDA) method (developed in Turner & Sederberg, 2014a; Holmes, 2015 and applied in Holmes et al., 2016; Holmes & Trueblood, 2017; Trueblood et al., 2018; Miletić, Turner, Forstmann, & van Maanen, 2017) to numerically approximate model log likelihoods for use in the Bayesian parameter estimation process.

Results (Figures 3) show that the type of experimental design greatly influences the ability to estimate these cognitive parameters (see Figure 4 for recovery results for other parameters). When evidence is fixed, parameter recovery is not possible on the basis of choice and RT data alone, and it appears to be essentially impossible to distinguish between integration versus urgency dominated strategies. When evidence changes over time, however, estimates are much more precise, and although appreciable uncertainty remains, high, medium, and low urgency or leakage can be distinguished. There is one caveat to this. Leakage is only identifiable when the leakage rate is above *L* = 1. This is to be expected. Recall that the timescale associated with evidence leakage is 1/*L*. Thus low values of *L* are associated with long time constants (greater than 1 second). If the time constant associated with leakage is longer than the typical length of a trial, for all intents and purposes leakage would be approximately absent and therefore not identifiable. Thus, for the physiologically relevant range of leakage rates, all parameters are identifiable when changing information is utilized.

**Figure 3.**
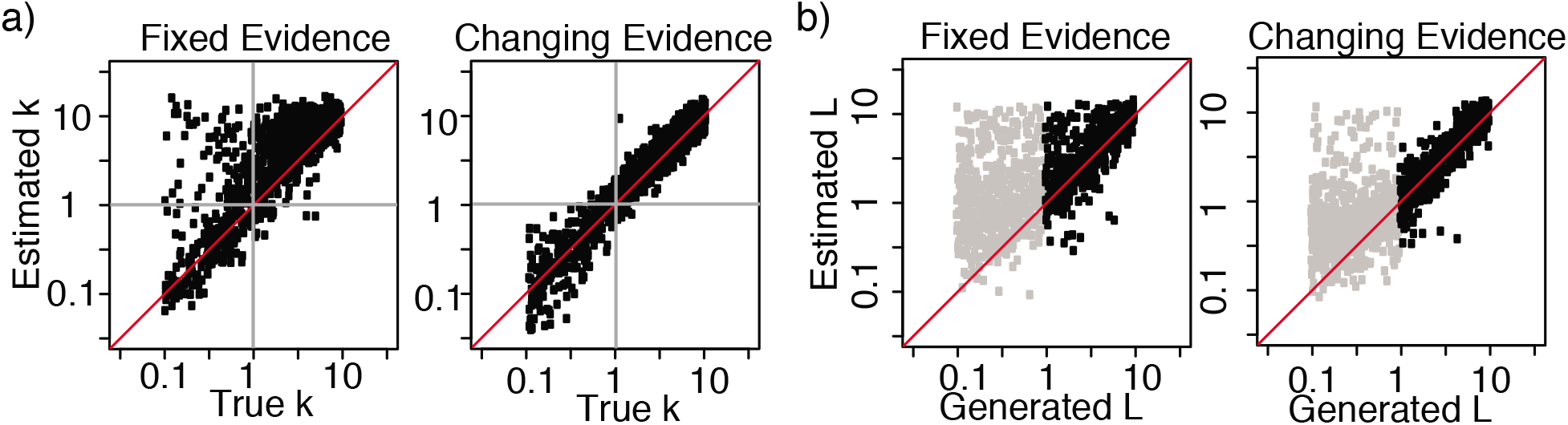
Specificity of urgency ratio and leakage constrained by RT data. Synthetic data sets were simulated out of the generalized UGM model for varying values of four parameters (*k, L, a, v*) and the model was fit using Bayesian methods to the resulting synthetic data. Results show the estimated (mean of the posterior) value of *k* and *L* versus the true value of *k* and *L* used to generate the synthetic data. Each point represents a single (true, estimated) value of the urgency ratio and the axes are presented on a log scale with the gray vertical and horizontal lines indicating the critical value of *k* = 1 and *L* = 1. The red line indicates the line of agreement, true value = estimated value. The first and third panels depict the quality of recovery when fixed, unchanging evidence is used. The second and fourth panels depict the quality when changing information is used. In the leakage recovery panels, gray dots are those with leakage values *L* < 1, which correspond to scenarios where leakage would be of limited importance on the typical 1-2 sec speeded decision timescale.

**Figure 4.**
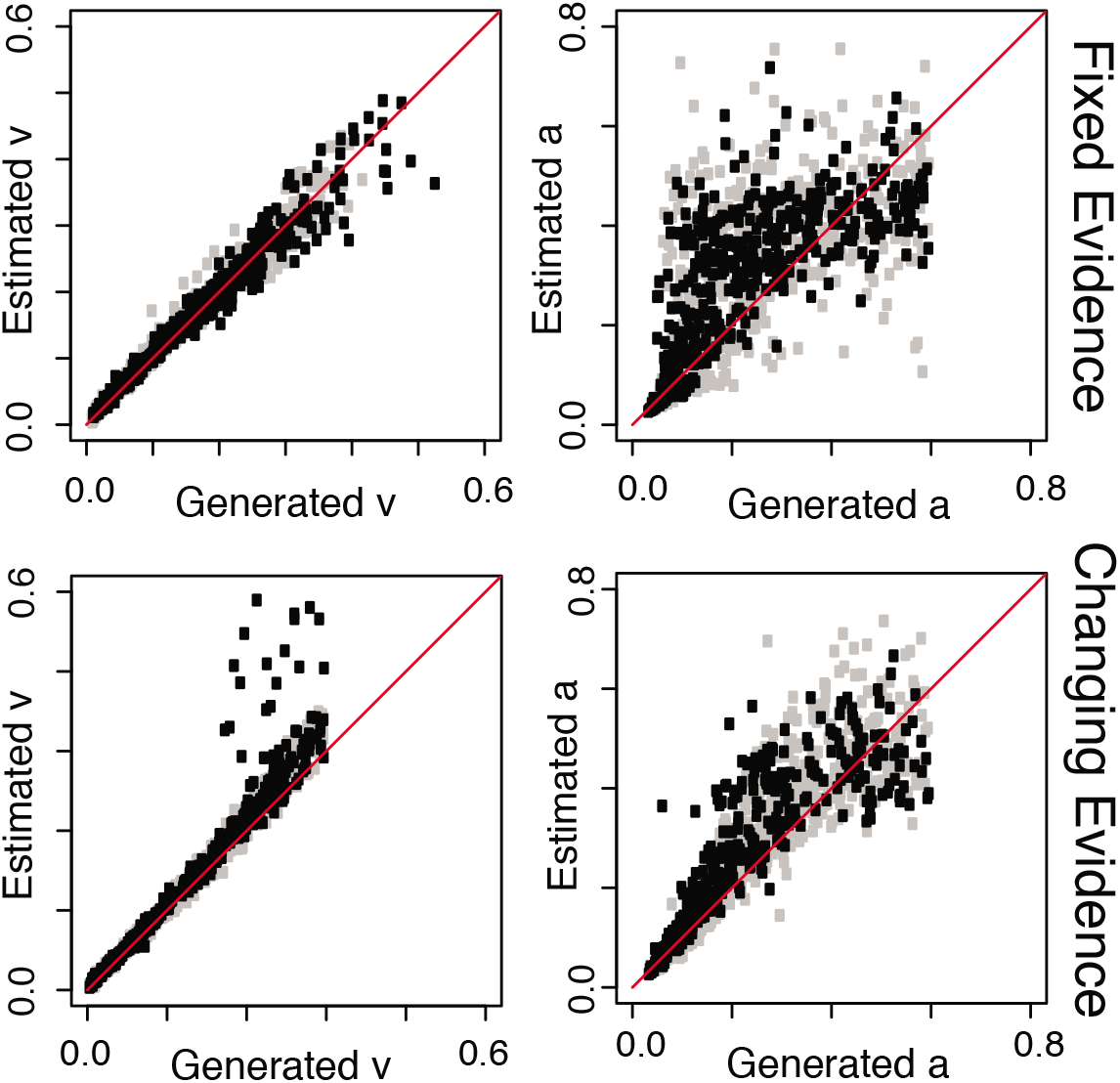
Specificity of drift and threshold constrained by RT data. Synthetic data sets were simulated out of the generalized UGM model for varying values of four parameters (*k, L, a, v*) and the model was fit using Bayesian methods to the resulting synthetic data. Results show the quality of recovery for the two parameters (*v, a*) when fixed (top) and changing (bottom) evidence are used. Gray dots are those with leakage values *L* < 1, which correspond to scenarios where leakage would be of limited importance on the typical 1-2 sec speeded decision timescale.

In conclusion, it is crucially important when investigating the role of urgency or evidence leakage to consider experimental paradigms where information changes over time. In the absence of such changes, it is difficult to constrain these cognitive parameters with choice and RT data alone. When a richer experimental paradigm utilizing changing information is used however, this generalized model has the potential to be used as an tool to estimate the relative strength (or even presence / absence) of these factors. In Experiments 1-4, we design and utilize a changing information paradigm to facilitate the estimation of these parameters in human participants.

### Experiment 1

Our experiments involve perceptual decisions about dynamic stimuli where the properties of the stimuli change partway through the trial, similar to Holmes et al. (2016). Whereas most past research on perceptual decision-making has used “stationary” decision tasks, where information is fixed or changes randomly around a central tendency (e.g., classic Random Dot Motion task), our paradigm switches the perceptual evidence from favouring one choice to another during the course of deliberation. As discussed above, this type of changing information allows for improved estimation of key cognitive parameters in the gUGM.

In the first experiment, we varied the strength of information before and after the change of information. On some trials, strong early (pre-switch) information is followed by weak late (post-switch) information. On other trials, weak pre-switch information is followed by strong post-switch information. Thus, there is an asymmetry in information strength, and consequentially difficulty, for early and late information. In addition to investigating the degree to which urgency and evidence leakage are present in this task, we will also examine the impact of this strength asymmetry on drift rates in the model to determine whether participants’ perception of evidence strength is veridical or whether one piece of information influences the perception of the next.

## Method

### Participants

A total of 34 Vanderbilt University students participated for course credit. All participants gave informed consent to participate. We set a sample size of about 30 participants in each experiment with accuracy of at least 70% on the practice trials. We continued to collect data until we met our sample size goal. This experiment and all remaining experiments were approved by the Vanderbilt University Institutional Review Board.

### Stimuli

Stimuli were ‘flashing grids’, a square grid filled with two different colored elements (see example in Figure 5). The two colors were orange (rgb: 251, 209, 132) and blue (rgb: 13, 255, 255). The grid flickered so that every three frames the elements were randomly rearranged. Each element in the grid was a 10×10 pixel square. The grid had 20 elements in each row and 20 elements in each column for a total of 400 colored elements. The grids were presented on a gray background (rgb: 95, 95, 95) and displayed on a 60 Hz monitor. In some of the trials, the proportion of blue to orange elements remained constant throughout the trial. In the remaining trials, the proportion of blue to orange elements changed once part way through the trial. On these ‘switch’ trials, the largest proportion color at the start of the trial became the smallest proportion color after the switch (e.g., if 60% of the elements were blue at the start of the trial, then it might switched to 60% orange).

**Figure 5.**
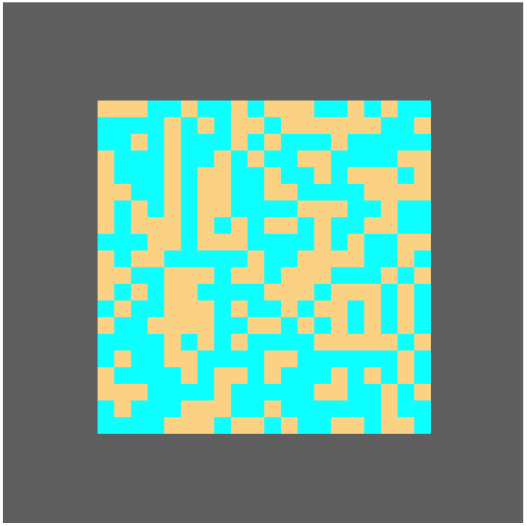
Flashing Grid Stimulus. Each grid was filled with elements of two colors (blue and orange) and presented on a gray background. The location of the elements was randomly rearranged every three frames. Participants were asked to decide if the grid had more blue or orange elements. On some trials, the proportion of colored elements changed part way through the trial.

### Procedure

At the beginning of the experiment, participants were instructed that they would see a flashing grid filled with blue and orange elements and their task was to decide if there are more blue or orange elements in the grid. They were told “you will enter your choices by pressing the ‘z’ key if you think the grid has more orange elements and ‘m’ key if you think the grid has more blue elements.” Participants were asked to place the index finger of their left hand of the ‘z’ key and the index finger of their right hand on the ‘m’ key throughout the experiment. They were also told that they would complete many blocks of trials, that some trials would be harder than others and that they would sometimes receive feedback about their responses. Participants were asked to respond to the best of their ability as quickly as possible, but were not told that the proportion of colored elements could possibly change during some trials.

At the beginning of each trial, they viewed a fixation cross for 250ms, then there was a blank screen for 100ms, followed by the stimulus. Participants had up to 2s to view the stimulus and give a response or the trial terminated by itself and a non-response was recorded. Otherwise, the trial terminated immediately after a response was made. The fixation cross for the next trial appeared immediately after the termination of the previous trial.

Participants completed 21 blocks of trials. Blocks 1-4 only contained ‘non-switch’ trials where the proportion of elements remained constant throughout the trial. The first block contained 20 practice trials where 75% of the elements in the grid were the same color (in 10 trials 75% of the elements were orange and in 10 trials 75% of the elements were blue). Participants received trial-by-trial feedback on their responses in this block. This feedback was displayed for 500 ms, and consisted of “correct”, “wrong”, or “invalid response” for invalid key presses. Feedback was followed by a blank screen for 250ms. In blocks 2-19, participants did not receive trial-by-trial feedback. However, they did receive block feedback. At the end of each block, participants were told their accuracy on that block as well as their average response time. The second block contained 72 additional practice trials where 57% of the elements in the grid were the same color. In blocks 3 and 4, participants completed 72 practice trials where 53% of the elements in the grid were the same color.

Blocks 5-19 were the main experiment and consisted of 72 trials each, where half were nonswitch trials and half were switch trials. The 36 non-switch trials were divided so that 18 trials had strong evidence (57% of the elements in the grid were the same color) and 18 trials had weak evidence (53% of the elements in the grid were the same color). The 36 switch trials in each block were divided so that 18 trials had strong pre-switch evidence followed by weak post-switch evidence. In these trials, 57% of the elements in the grid were the same color before the switch and 53% of the elements in the grid were the other color after the switch. For example, if 57% of the elements were orange before the switch, then 53% of the elements were blue (47% orange) after the switch. The remaining 18 trials had weak pre-switch evidence followed by strong post-switch evidence. In these trials, 53% of the elements in the grid were the same color before the switch and 57% of the elements in the grid were the other color after the switch. For example, if 53% of the elements were orange before the switch, then 57% of the elements were blue (43% orange) after the switch. The switch time was fixed to 350 ms for all switch trials. The trials in each block were randomly presented.

Blocks 20 and 21 tested whether participants could detect changes in color proportions. During these blocks, they were instructed to only respond if the proportion of colored elements changed. They were told to withhold responses if the proportion remained constant. Block 20 contained 24 practice trials with feedback. At the end of each trial, participants saw a message “correct”, “wrong”, or “invalid response” displayed for 500ms. Block 21 contained 72 trials with no feedback. In both blocks, half of the trials were switch trials and the other half were non-switch trials, randomly ordered. Like blocks 5-19, there were two difficulty levels of non-switch trials and two types of switch trials.

## Behavioral Results

Our exclusion criterion was accuracy lower than 70% in block 2 (the practice trials with strong evidence). All participants met the accuracy criterion and thus none were removed from the analyses below. The choice data was analyzed using generalized linear mixed-effects models (GLMMs). In this experiment, there were two main factors of interest - stimulus strength and response time quantile. The latter was determined on an individual basis by calculating nine evenly spaced RT quantiles and then calculating the choice proportions for each of the resulting 10 bins. For all of the GLMMs considered, we used a probit link function and allowed for random intercepts and slopes for each subject, following the recommendation of Barr, Levy, Scheepers, and Tily (2013) in using a maximal random effects structure. All GLMMs were fit using the fitglme function in Matlab.

First, we analyzed the choice data for the non-switch trials. We fit four different GLMMs, examining the inclusion of various fixed effects (i.e., a null model, a model with only stimulus strength, a model with only RT quantile, and a model with both stimulus strength and RT quantile). All models included an intercept. The model with both fixed effects of stimulus strength and RT quantile had the lowest BIC and AIC values. The best fitting model showed that accuracy differed significantly by stimulus strength (*β* = 0.227, t(676) = 3.68, p < 0.01, 95% CI [0.11, 0.35]), suggesting that participants were more accurate on trials with strong stimulus information (see Figure 7a). There was no main effect of RT quantile (*β* = −0.025, t(676) = −1.62, p = 0.106, 95% CI [−0.05, 0.01]). However, the interaction between stimulus strength and RT quantile was significant (*β* = 0.025, t(676) = 3.40, p < 0.01, 95% CI [0.01, 0.04]). As shown in Figure 7a, accuracy was highest for the middle RT quantiles, suggesting both fast and slow errors (Luce, 1986; Ratcliff & Rouder, 1998).

**Figure 6.**
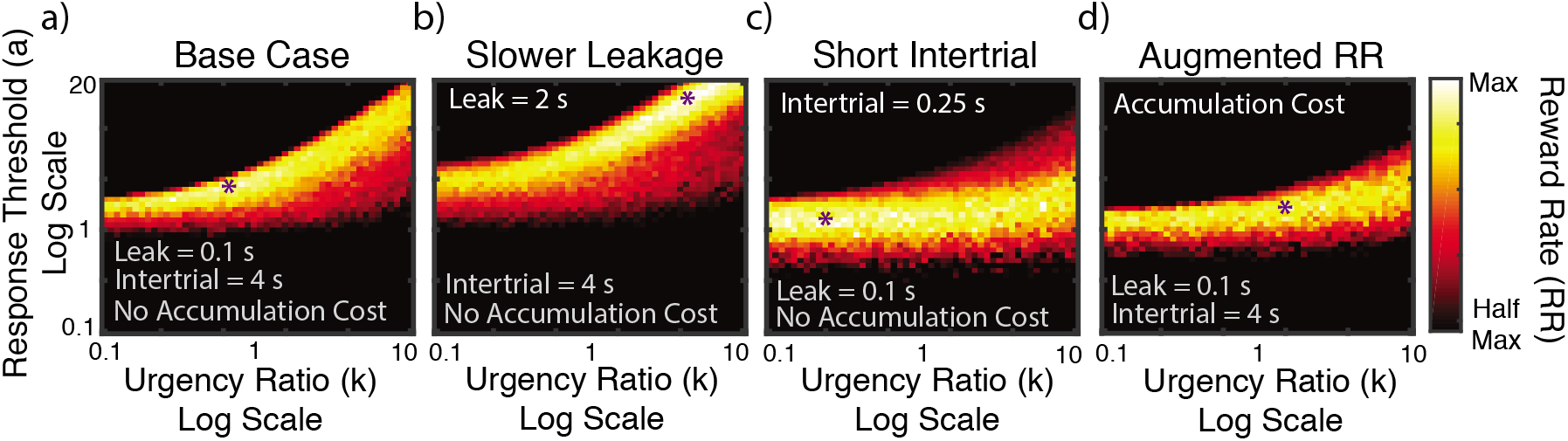
Influence of Urgency on Reward Rate. **Panels a-d)** These heatmaps show the reward rate (RR) as a function urgency ratio and response threshold in four scenarios. In the base scenario: *L* = 4, *t_s_* = 0.2, *t_I_* = 4, and the cost function is *C*(*t*) = 0. In the remaining plots (b-d), the relevant parameter that changes is indicated in the top left hand corner. In panel (d), the cost function takes the form *C*(*t*) = 1/2*t*^2^, representing a linearly increasing cost per unit time. Purple stars in all cases indicate the maximal reward rate for that plot and in all cases, black regions indicate those parameter sets where *RR* < 0.5 * *RR_max_*.

**Figure 7.**
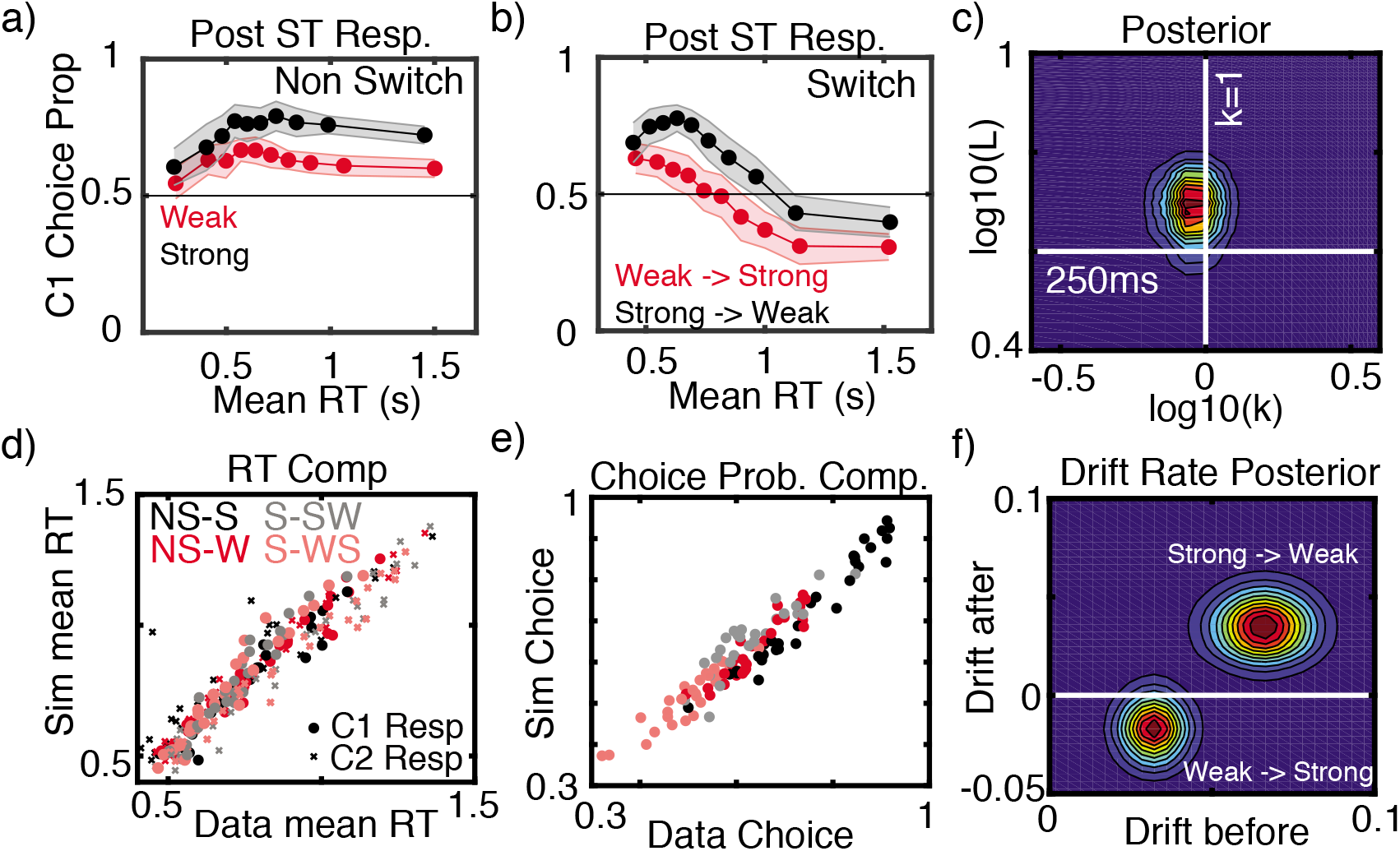
Experiment 1 Results. **Panel a)** Mean choice proportions for the majority color on non-switch trials with weak (red) and strong (black) information. Choices are plotted for nine evenly space RT quantiles. Error band shows the 95% confidence interval. **Panel b)** Mean choice proportions for the first majority color on switch trials with responses after the switch time of 350 ms. Trials with weak information after the switch (i.e., strong information before the switch) are in black and trials with strong information after the switch (i.e., weak information before the switch) are in red. Choices are plotted for nine evenly space RT quantiles. Error band shows the 95% confidence interval. **Panels d-e)** Demonstration of quality of model fit. Each dot on both panels depicts the true and simulated (from mean parameters) mean response time and choice proportion for a single participant in a single experimental condition. The legend abbreviations correspond to the following. NS and S refer to non switch and switch trials respectively while W and S correspond to weak and strong stimuli respectively. Thus S-WS corresponds to a switch trial for which the stimulus changes from weak to strong. In (d), C1 (resp. C2) indicates the mean RT for responses for the first color (second color resp.). Data that lie on the y=x (predicted = true) line indicate the model is providing a good fit to data. **Panel c)** Joint posterior distribution for the hyper means for the urgency ratio (*k*) and leakage (*L*) parameters. **Panel f)** Joint posterior distribution for the hyper means for the drift rates before and after the switch in both switch conditions. Two joint posteriors are plotted. In one, the weak drift before is jointly plotted with the strong drift after to correspond to the weak → strong condition. In the other, the strong drift before is plotted against the weak drift after. The white line delineates positive from negative drift rates after the switch. A positive value of the drift indicates accumulation for the first color shown and a negative drift for the second color shown.

Next, we analyzed the switch trials with responses after the switch time of 350 ms (that is, we excluded switch trials with responses before the switch time). For this analysis, we examined choices for the first majority color (i.e., the color with the most elements prior to the switch). Since this analysis only uses trials with responses after the switch, choices for the first majority color can be considered incorrect. Similar to the non-switch trial analysis, we fit four different GLMMs. As in that analysis, the model with both fixed effects of stimulus strength and RT quantile had the lowest BIC and AIC values. The best fitting model showed that choices differed significantly by stimulus strength (*β* = 0.525, t(676) = 7.16, p < 0.01, 95% CI [0.38, 0.67]) as well as RT quantile (*β* = −0.11, t(676) = −5.18, p < 0.01, 95% CI [−0.15, −0.07]). The interaction was not significant (β = −0.01, t(676) = −0.75, p = 0.452, 95% CI [−0.02, 0.01]). As shown in Figure 7b, choices for the first majority color decreased with RT quantile. That is, slower responses tended to result in more choices for the second majority color (i.e., the “correct” answer after the switch). The figure also shows that there were fewer choices for the first majority color in the weak to strong trials (e.g., a trial going from 53% orange to 57% blue) as compared to the strong to weak trials. In other words, when the second stimulus information was strong, participants more often selected the second majority color (i.e., the correct response).

We also examined detection ability using the results from Block 21. The mean hit rate was 0.56 and the mean false alarm rate was 0.49. Discriminability (average d’ = .20) was significantly above zero, implying that participants could detect changes in this task (t(33) = 2.63, p = 0.013).

## Modeling Results

Hierarchical Bayesian fitting of the gUGM to the full choice-RT distributions for this data demonstrates three important points. Figure 7c shows the joint hyper level posterior for the urgency ratio and leakage rate. These results demonstrate that the urgency ratio is at moderate levels (*k* ~ 1) while the leakage rate is relatively quick (the leakage timescale is faster than 250ms). Thus time varying urgency and leakage both appear to be present here. We note that the leakage timescale is similar to that proposed in (Cisek et al., 2009). Critically however, we have estimated this directly from human data here rather than simply fixing it. The estimated value of the urgency ratio *k* ~ 1 is however on the low end of the spectrum that was estimated in (Thura et al., 2014) where results indicated this ratio should fall in the *k* ∈ [0.5, ∞] range.

The third observation relates to how the initially presented information influences the perception of subsequent stimulus information after the change. If the decision process were completely veridical, one would expect the perception of the second stimulus to be independent of what precedes it. Rather than build this assumption into the model, we estimated separate drift rates for the first and second stimulus segments in both the strong → weak and weak → strong conditions. Figure 7f shows the joint posteriors for the before and after drift rates for the two conditions separately (the two peaks in this figure are two separate posteriors). The horizontal axis depicts the drift rate before the switch and the vertical after the switch. For both the before and after drift rate axes, a positive value indicates drift favor of the first stimulus while a negative value indicates drift in favor of the second stimulus.

Results show that in both conditions, the first stimulus affects the perception of the second. If we consider only the before switch drift rates, as expected, the drift rate corresponding to the weak stimulus is lower than that of the strong stimulus. The posterior mean drift rates for the weak and strong stimuli are ~ 0.03 and 0.06 respectively. If the decision process is veridical, it would be expected that in the weak-to-strong case, the drift rate would transition from 0.03 → –0.06 with the negative sign indicating a flip in drift for the second stimulus. Comparison of the pre and post-switch drifts in the weak-to-strong condition shows that this is not the case. While the drift does indeed change sign as expected, it transitions from 0.03 → –0.02, indicating that the estimated drift is substantially attenuated (–0.02 as compared to the –0.06 veridical). The strong-to-weak condition shows an even more substantial effect. In this case the post-switch drift rate is still positive, indicating drift in favor of the first stimulus (which is objectively incorrect after the change in information).

These results are consistent with the hypothesis that pre-switch information attenuates the perception of post-switch information. When strong follows weak, the post-switch drift rate indicates drift for the second stimulus, but with an attenuated magnitude. When weak follows strong, the initial stimulus overwhelms the subsequent stimulus and the post-switch drift continues to support the first stimulus.

For comparison, we also fit two restrictions of the full gUGM model. The first restriction fixed the urgency ratio at *k* = 0, which leads to a model that does not encode time varying urgency (e.g., a pure leaky integrator). The second restriction considers the *k* → ∞, which corresponds to the standard UGM model that has most commonly been studied. Table 2 demonstrates that both restricted models are > 300 points worse according to DIC.

**Table 2.** Table of DIC differences between the full gUGM (unrestricted and estimated urgency ratio k) and two limiting simplifications of the full gUGM. Values indicate the difference in DIC values between the full gUGM and the model being compared to it (thus no value is given for the full gUGM). When comparing the full model to the standard UGM on the Experiment 1 data set, the DIC value for the standard UGM is 865 points higher than for the full gUGM, indicating the standard UGM provides a much worse fit to data in that case. Since Experiments 3 and 4 are factorial combinations of 1 and 2, we only perform this comparison for the two base experimental designs.

| Model | Exp. 1 | Exp. 2 |
| --- | --- | --- |
| Full gUGM | - | - |
| No Urgency ($k=0$) | 358 | 3 |
| Standard UGM ($k=\infty$) | 865 | 86 |

Note that we are not advocating for the use of this model comparison approach as the primary means of drawing conclusions in this case. One of the benefits of the gUGM model is that it can be used for parameter inference, allowing one to estimate the urgency parameter. If the *k* = 0 or *k* = ∞ limits lead to better accounting of the data, the model can converge to those limits. These comparison results do however further support our parameter based inference that a balance between integration and urgency is required to account for our data, and models that do not allow for this possibility perform much worse.

## Conclusions

In conclusion, the behavioral and modeling results demonstrate that 1) participants’ urgency varies over time, although not as much as has been reported previously, 2) evidence leakage is present with a timescale of 200-250ms, consistent with some previous estimates, and 3) information presented early in the trial substantially influences the perception of subsequent information, consistent with the results of Holmes et al. (2016).

### Experiment 2

Experiment 1 shows that the strength of early information impacts decisions about later information. In that experiment, the duration of early information was fixed (at 350 ms) for all trials. It is possible that the duration of early information (even if the strength of the information is held constant) also impacts the perception of subsequent information. In Experiment 2, we varied the switch time across trials with symmetric changes in the strength of information. We hypothesize that longer pre-switch durations will have a great impact on the perception of postswitch information than shorter pre-switch durations. As in Experiment 1, our goal is to measure three important aspects of the decision process: urgency, evidence leakage, and drift rates.

## Method

### Participants

A total of 51 Vanderbilt University students participated for course credit. All participants gave informed consent to participate. Similar to Experiment 1, we set a sample size of about 30 participants with accuracy of at least 70% on the practice trials. We continued to collect data until we met our sample size goal.

### Stimuli

The stimuli and instructions were the same as Experiment 1.

### Procedure

Participants completed 24 blocks of trials. Blocks 1-7 only contained ‘non-switch’ trials where the proportion of elements remained constant throughout the trial. Similar to the Experiment 1, the first block contained 20 practice trials where 75% of the elements in the grid were the same color (in 10 trials 75% of the elements were orange and in 10 trials 75% of the elements were blue). Again, participants received trial-by-trial feedback on their responses in this block. In blocks 2-22, participants received block feedback, but not trial-by-trial feedback. The second block contained 72 additional practice trials where 54% of the elements in the grid were the same color. In the third block, participants completed 72 trials, which were composed of 24 trials at three different difficulty levels: 52%, 54%, or 56% of the elements had the same color. At the conclusion of the third block, one of the three difficulty levels was selected for the remainder of the experiment using the following algorithm to account for individual differences in ability so that performance was away from floor and ceiling:

1. If the participant’s accuracy was exactly 75% for a specific difficulty level, then this level was selected. If more than one difficulty level achieved 75% accuracy, then the hardest level was selected.
2. If no difficulty level achieved accuracy of exactly 75%, then the level with accuracy higher and closest to 75% was selected. If there was a tie between difficulty levels, then the hardest one was selected.
3. If none of the difficulty levels achieved 75% accuracy, then the easiest level (i.e., 56%) was selected.

Blocks 4-7 each contained 72 trials with the difficulty level selected at the end of block 3. These four blocks provided the participant with additional practice at the selected difficulty level and helped control for learning effects before the main part of the experiment.

Blocks 8-22 were the main experiment and consisted of 72 trials each, where half were nonswitch trials and half were switch trials. The 36 switch trials in each block were divided so that 12 trials had a switch time of 150 ms, 12 trials had a switch time of 250 ms, and the remaining 12 trials had a switch time of 350 ms. The trials in each block were presented in a random order.

Blocks 23 and 24 were similar to blocks 20 and 21 in Experiment 1. These blocks tested whether participants could detect changes in color proportions. All three switch times were used and the difficulty level for these blocks was the same as in the main task.

## Behavioral Results

Participants were excluded if their average accuracy in blocks 4-7 was less than 70%. This left 30 participants for data analyses.^1^ Similar to Experiment 1, the choice data was analyzed using GLMMs. In this experiment, there were two main factors of interest - switch time and RT quantile, where the latter was determined as in Experiment 1. As before, we assumed by-subject random intercepts and slopes. Since there was only one type of non-switch trial (i.e., the proportion of colors was the same for all non-switch trials for an individual), we only analyzed the switch trials.

For the analysis of the switch trials, we examined choices for the first majority color in trials with responses after the switch time, similar to Experiment 1. We fit four different GLMMs, examining the inclusion of various fixed effects (i.e., a null model, a model with only switch time, a model with only RT quantile, and a model with both switch time and RT quantile). All models included an intercept. The model with both fixed effects of switch time and RT quantile had the lowest BIC and AIC values. The best fitting model showed that choice differed significantly by switch time (*β* = 4.93, t(896) = 13.16, p < 0.01, 95% CI [4.19, 5.66]) as well as RT quantile (*β* = −0.04, t(896) = −2.75, p < 0.01, 95% CI [−0.08, −0.01]). The interaction was also significant (*β* = −0.36, t(896) = −7.53, p < 0.01, 95% CI [−0.45, −0.27]). As shown in Figure 9a, choices for the first majority color decreased with RT quantile. That is, slower responses tended to result in more choices for the second majority color, similar to Experiment 1. The figure also shows that there were fewer choices for the first majority color in trials with the earliest switch time (150 ms switch) as compared to the trials with later switch times (250 and 350 ms). In other words, when the duration of the first stimulus was short, participants more often selected the second majority color.

We also examined detection ability using the results from Block 24. The mean hit rate was 0.53 and the mean false alarm rate was 0.46. Similar to Experiment 1, discriminability (average d’ = .17) was significantly above zero, implying that participants could detect the changes of information (t(29) = 3.83, p < .001).

## Modeling Results

Based on the results of Experiment 1, Experiment 2 was designed to test how the duration of the pre-switch stimulus influences decisions. Our hypothesis was that the duration itself should not substantially alter the leakage or urgency characteristics of the decision process. However, if early information attenuates the perception of late information, we would naturally expect longer durations of the initial stimulus phase to have a larger effect.

To test this, we fit the gUGM allowing different pre and post-switch drift rates for each of the three duration conditions (quality of model fit shown in Figure 8). Results largely confirmed our hypothesis. Figure 9b demonstrates that the urgency ratio takes on a mean value of ~ 1.6 while the leakage timescale is again ~ 250ms. Analysis of the drift rates (Figure 9 (c-e)) further shows two basic results. First, even a quick 150ms initial stimulus duration affects the post switch drifts. Second, that effect becomes stronger as the duration of the first stimulus presentation increases.

**Figure 8.**
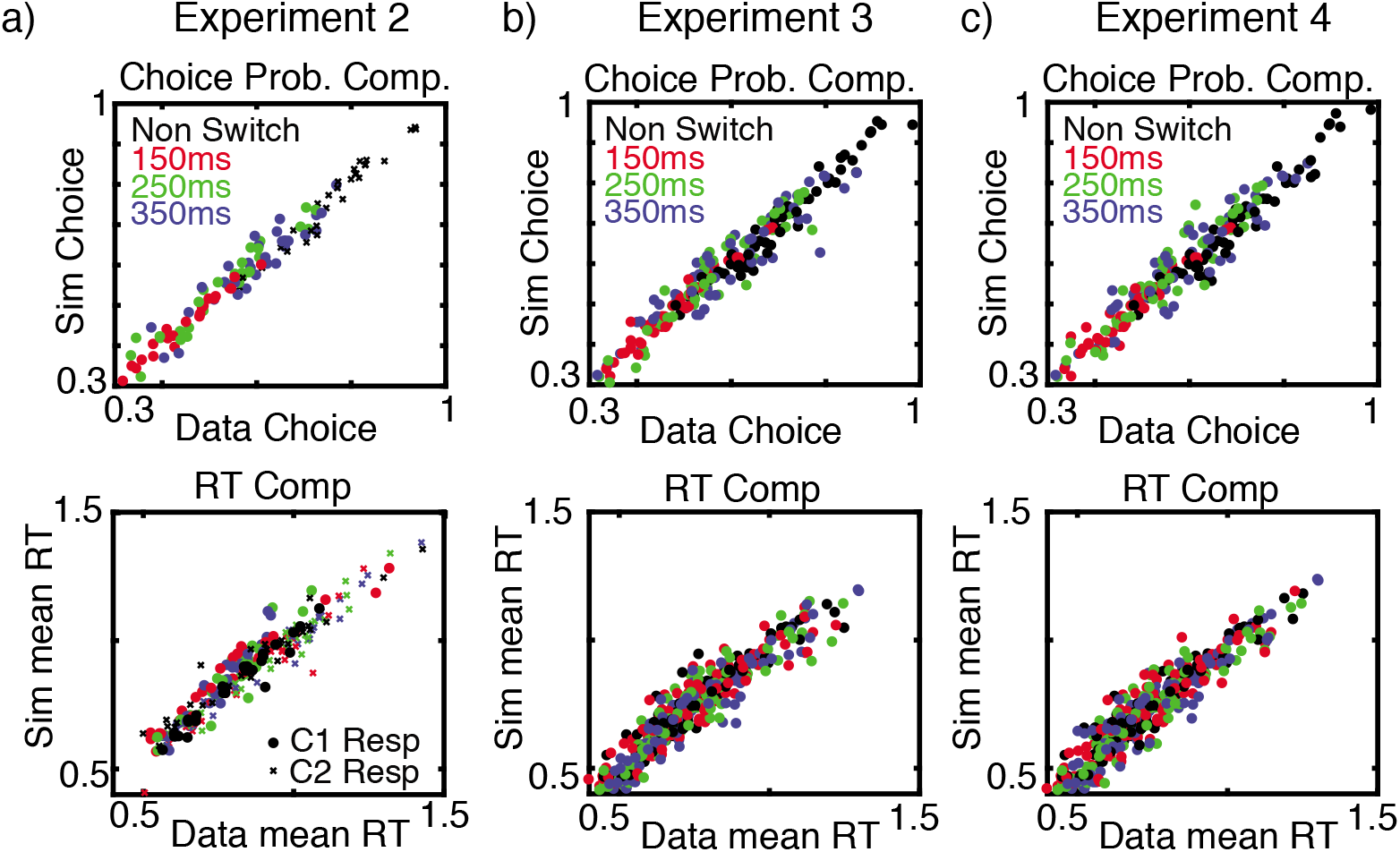
Quality of Model Fit to Experiments 2-4. Quality of model fit. Each dot on both panels depicts the true and simulated (from mean parameters) mean response time and choice proportion for a single participant in a single experimental condition.

**Figure 9.**
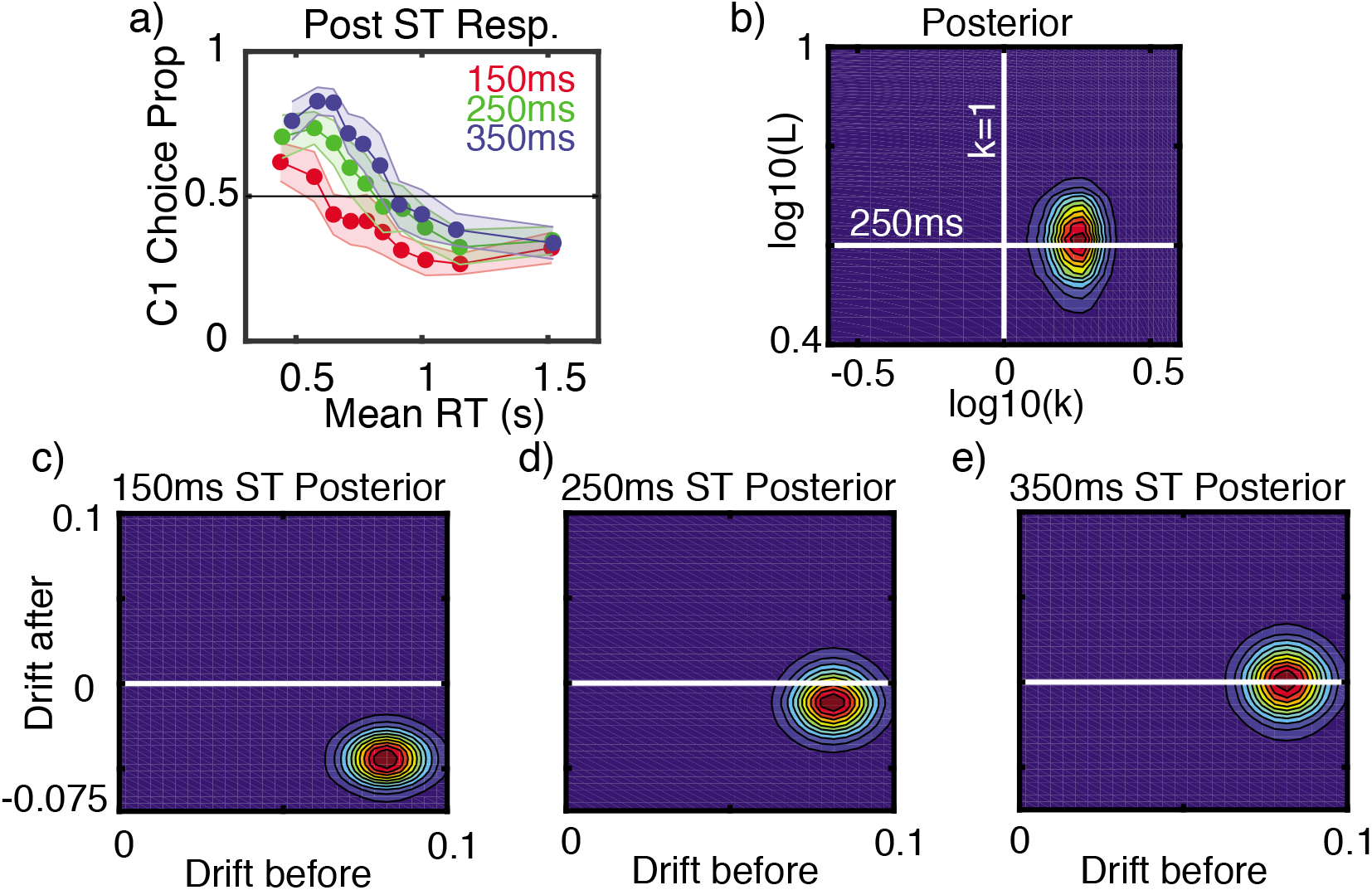
Experiment 2 Results. **Panel a)** Mean choice proportions for the first majority color on switch trials with responses after the switch time. Choices are plotted for nine evenly space RT quantiles. Error band shows the 95% confidence interval. **Panel b)** Joint posterior distribution for the hyper means for the urgency ratio (*k*) and leakage (*L*) parameters. **Panesl c-e)** Joint posterior distribution for the hyper means for the drift rates before and after the switch for the three switch conditions. Plotting conventions are the same as in Figure 7.

Table 2 shows once again that the standard UGM (*k* = ∞) provides a significantly worse accounting of the experimental data when compared to the full gUGM (according to DIC). The *k* = 0 limit on the other hand performs comparably to the full gUGM. We note once again that in Experiment 1, the *k* = 0 leaky integrator limit performed vastly worse than the full gUGM. Thus the preponderance of evidence still supports the conclusion that a balance between urgency and integration best accounts for observations.

## Conclusions

These results once again demonstrate that the decision process involves a moderate increase in urgency over time, evidence leakage, and that the pre-switch stimulus influences the perception of subsequent stimuli in a duration dependent way, where a longer duration yields a larger influence on subsequent drift rates.

### Experiment 3

Experiments 1 and 2 examined the impact of strength and duration of information separately. In Experiment 3, we combined these two factors and examine the influence of both information strength and switch time on people’s decisions about changing information. In this experiment, we manipulated the strength of the pre and post-switch information similarly to Experiment 1 and the switch time similarly to Experiment 2.

## Method

### Participants

A total of 35 Vanderbilt University students participated for course credit. All participants gave informed consent to participate. Similarly to the other experiments, we set a sample size of about 30 participants in each experiment with accuracy of at least 70% on the practice trials. We continued to collect data until we meet our sample size goal.

### Stimuli

The stimuli and instructions were the same as in Experiments 1 and 2.

### Procedure

The procedures for this experiment were very similar to Experiment 1. The practice trials in Blocks 1-4 were identical to those in Experiment 1. The main part of the experiment consisted of blocks 5-19 with 72 trials per block. The 36 non-switch trials in each block where divided into two difficulty levels as in Experiment 1. The 36 switch trials in each block were subdivided into 6 types by crossing three switch times (150 ms, 250 ms, and 350 ms, as in Experiment 2) with two pairs of pre and post switch information strengths (strong evidence followed by weak evidence and vice versa, as in Experiment 1). Blocks 20 and 21 were similar to Experiment 1, except for the inclusion of the three different switch times on switch trials.

## Behavioral Results

Similar to Experiment 1, participants were excluded if their average accuracy in block 2 was less than 70%. This left 29 participants for data analyses. As in the previous experiments, the choice data was analyzed using GLMMs. In this experiment, there were three main factors of interest - stimulus strength, switch time, and RT quantile. As before, we assumed by-subject random intercepts and slopes.

First, we analyzed the choice data for the non-switch trials (see Figure 10a). We fit four different GLMMs, examining the inclusion of various fixed effects (i.e., a null model, a model with only stimulus strength, a model with only RT quantile, and a model with both stimulus strength and RT quantile). Note that switch time is not included in these models because it is irrelevant for the non-switch trials. All models included an intercept. The model with both fixed effects of stimulus strength and RT quantile had the lowest BIC and AIC values. The best fitting model showed that accuracy differed significantly by stimulus strength (*β* = 0.273, t(576) = 4.12, p < 0.01, 95% CI [0.14, 0.40]), showing that participants were more accurate on trials with strong stimulus information. There was no effect of RT quantile (*β* = −0.029, t(576) = −1.78, p = 0.075, 95% CI [−0.061, 0.003]). However, the interaction was significant (β = 0.020, t(576) = 2.53, p = 0.01, 95% CI [0.004, 0.035]). These results are consistent with the findings from Experiment 1.

**Figure 10.**
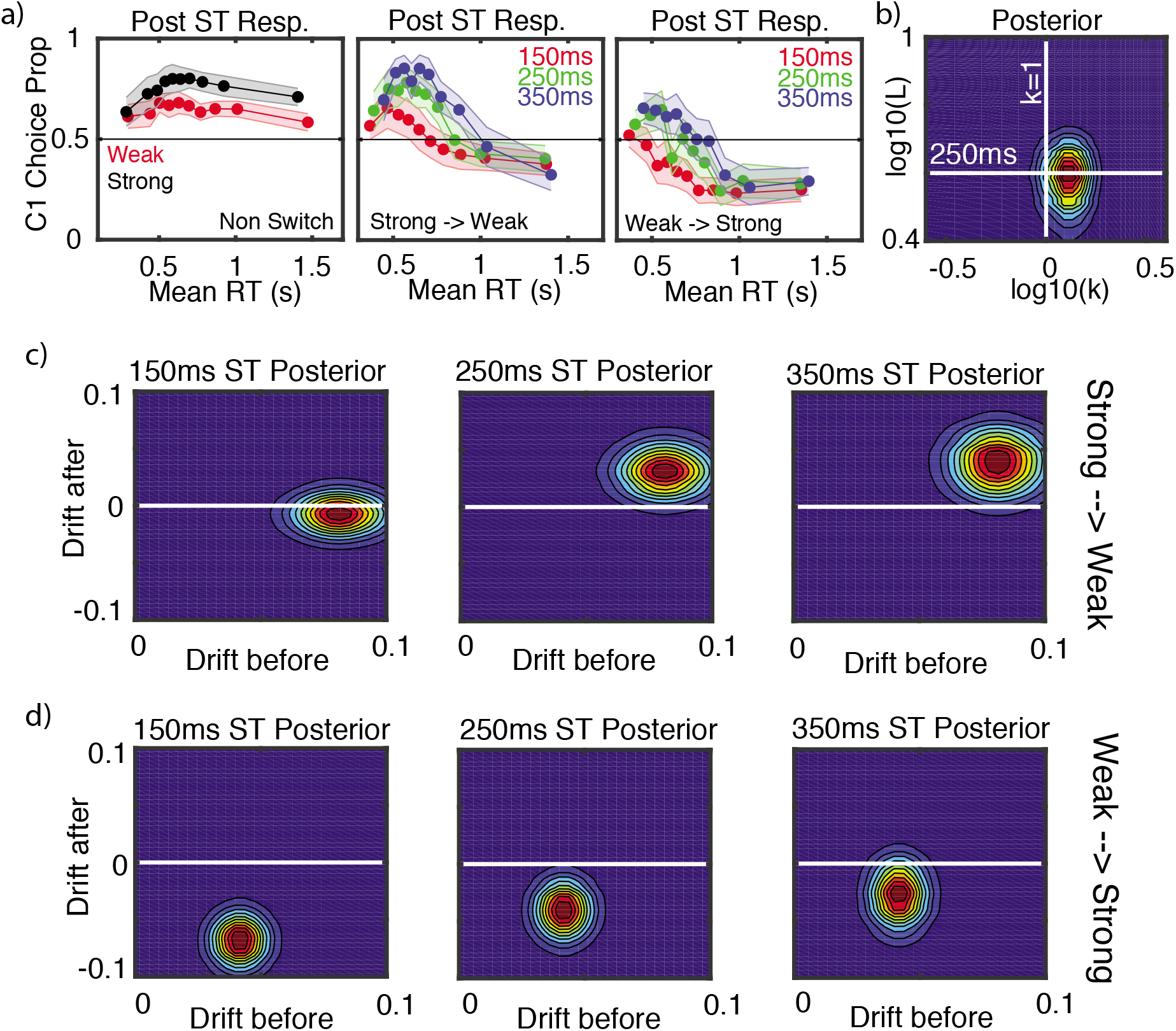
Experiment 3 Results. **Panel a)** Mean choice proportions for the first majority color on non-switch trials (left), switch trials with responses after the switch time with strong followed by weak information (middle), and switch trials with weak followed by strong information (right). Choices are plotted for nine evenly space RT quantiles. Error band shows the 95% confidence interval.**Panel b)** Joint posterior distribution for the hyper means for the urgency ratio (*k*) and leakage (*L*) parameters. **Panesl c-d)** Joint posterior distribution for the hyper means for the drift rates before and after the switch for the six switch conditions (three switch times along two different strength conditions). Plotting conventions are the same as in Figure 7.

For the analysis of the switch trials, we examined choices for the first majority color in trials with responses after the switch time, similar to Experiments 1 and 2. We fit eight different GLMMs, examining the inclusion of various fixed effects (i.e., a null model, models with only a single factor, models with two factors, and a model with all three factors). All models included an intercept. The model with all three factors (stimulus strength, switch time, and RT quantile) had the lowest BIC and AIC values. The best fitting model showed that choices differed significantly by RT quantile (*β* = −0.13, t(1732) = −3.08, p < 0.01, 95% CI [−0.21, −0.05]). The fixed effects of switch time and stimulus strength were not significant. However, their interaction was significant (*β* = 1.51, t(1732) = 2.44, p = 0.015, 95% CI [0.29, 2.72]). The interaction of stimulus strength and RT quantile was also significant (*β* = 0.05, t(1732) = 2.09, p = 0.037, 95% CI [0.003, 0.103]). None of the other interactions were significant. As shown in Figure 10a, choices for the first majority color decreased with RT quantile. In addition, the effects of stimulus strength and switch times are similar to those seen in Figures 7b and 9a for Experiments 1 and 2.

We also examined detection ability using the results from Block 21. The mean hit rate was 0.56 and the mean false alarm rate was 0.45. As in Experiments 1 and 2, discriminability (average d’ = .29) was significantly above zero, implying that participants could detect changes in this experiment (t(28) = 3.77, p < .001).

## Modeling Results

Data from this experiment were fit with an extension of the model that allowed for different pre and post-switch drift rates in each of the six experimental conditions (quality of model fit shown in Figure 8). Results (Figure 10) are consistent with prior modeling results. The urgency ratio was estimatwed as *k* = 0.95 along with a leakage timescale of ~ 250ms, both of which are comparable to findings from Experiments 1 and 2. Further, the effect of initial stimulus information on post switch drifts follows the same patterns found in modeling results for Experiments 1, 2. That is, longer initial durations have more substantial effects on post switch drifts and stronger initial information has a stronger effect.

## Conclusions

The results of Experiment 3 are consistent with those found in the previous two experiments. That is, when we factorially combine the duration and strength manipulations from Experiments 1 and 2, we find similar results for urgency, evidence leakage, and drift rates.

### Experiment 4

In the previous experiments we examined how the duration and strength of early information impacts the processing of later information. In all of these experiments, the stimuli information was perceptual. In Experiment 4, we extend beyond perceptual tasks to ask how value impacts people’s ability to adapt to changed information. Decades of past research has shown a systematic asymmetry in choices for equivalent gains and losses (generally studied with monetary gambles, Tversky & Kahneman, 1992). For example, people generally attempt to avoid a loss of $5 more than seeking a gain of $5, a phenomenon known as loss aversion. Based on people’s general desire to avoid losses in combination with our results from Experiments 1-3, we predict that early negative information (i.e., a gamble with negative expected value at the start of a trial) will result in slower adaptation to later information than early positive information (i.e., a gamble with positive expected value at the start of a trial). We test this hypothesis by examining how people adapt to new information when a gamble changes from a negative to a positive expected value as compared to a gamble that changes from positive to negative expected value.

Experiment 4 uses the same flashing grid stimuli as Experiments 1-3; however, in this task the flashing grids represent gambles offering different probabilities of winning or losing amounts of money. Similar to Experiment 3, we manipulated the strength of pre and post-switch information and switch times.

#### Participants

A total of 30 Vanderbilt University students participated for course credit. All participants gave informed consent to participate. We set a sample size of 30 participants based on the previous experiments.

#### Stimuli

The stimuli were the same as in the previous experiments.

#### Procedure

The procedure for this experiment were very similar to Experiment 3. However, unlike the previous three experiments, participants were instructed that they would be making decisions about gambles. On each trial they had to decide to either accept or reject a gamble. The gamble was represented as a flashing grid with orange and blue elements. Participants were told that the color orange was associated with winning $5 and the color blue was associated with losing $5. They were also told that the proportion of orange elements represented their chance of winning $5 and the proportion of blue elements represented their chance of losing $5. Participants were also told that their goal was to earn as much money as possible (note that, in reality their choices were hypothetical and not actually paid). They were instructed to “enter your choice by pressing the ‘z’ key if you want to play the gamble and ‘m’ key if you do not want to play the gamble.” Participants were asked to place the index finger of their left hand of the ‘z’ key and the index finger of their right hand on the ‘m’ key throughout the experiment. Before beginning the first practice block, participants were reminded of the color / value associations (i.e., orange = win $5 and blue = lose $5) again.

Participants completed 21 blocks of trials. Blocks 1-4 only contained ‘non-switch’ trials where the proportion of elements remained constant throughout the trial. The first block contained 40 practice trials where 75% of the elements in the grid were the same color (in 20 trials 75% of the elements were orange, a positive expected value gamble, and in 20 trials 75% of the elements were blue, a negative expected value gamble), and participants received trial-by-trial feedback on their choices. To determine the amount earned on each trial, a preprogramed algorithm ran the gamble if selected and the results were recorded. If participants rejected the gamble or responded too late, they would earn $0 for that trial. When participants accepted the gamble and won, they received feedback stating, “Gamble Selected, Win $5”. When participants accepted the gamble and lost, they received feedback stating, “Gamble Selected, Lose $5”. If participants rejected the gamble, they received feedback stating “Gamble not Selected, Receive $0”. In blocks 2-19, participants did not receive trial-by-trial feedback. However, they did receive block feedback. At the end of each block, participants were told the amount they earned in that block, the total amount earned overall, and their average response time. Blocks 2-21 were identical to Experiment 3.

## Behavioral Results

Unlike the previous experiments, we did not exclude participants based on whether their average accuracy in block 2 was less than 70%. Since this experiment involves subjective risky decisions, accuracy is not well defined. However, we removed three participants from the analyses because their total earnings in the task were negative. This suggests that they were either not engaged in the task or did not understand the instructions. This left 27 participants for the data analysis and modeling. As in the previous experiments, the choice data was analyzed using GLMMs. In this experiment, there were four main factors of interest - gamble type (positive versus negative expected value), stimulus strength, switch time, and RT quantile. In this task, stimulus strength relates to the expected value (EV) of the gamble. “Strong” information implies that the gamble has a more extreme EV (further from zero) and “weak” information means that is has a less extreme EV (closer to zero). To retain consistency with Experiments 1-3, we will continue to use the “strong” and “weak” terminology. As before, we assumed by-subject random intercepts and slopes. In all of the analyses below, we coded responses according to the EV maximizing decision for the gamble. For positive EV gambles, the EV maximizing choice is ‘accept’. For negative EV gambles, the EV maximizing choice is ‘reject’.

First, we analyzed EV maximizing choices (i.e., accept for positive EV gambles and reject for negative EV gambles) on the non-switch trials. We fit eight different GLMMs, examining the inclusion of various fixed effects (i.e., a null model, models with only a single factor, models with two factors, and a model with three factors). Once again, switch time is not included in these models because it is irrelevant for the non-switch trials. All models included an intercept. The model with all three relevant factors (gamble type, stimulus strength, and RT quantile) had the lowest BIC and AIC values. The best fitting model showed that choices differed significantly by gamble type (*β* = 1.518, t(712) = 5.25, p < 0.01, 95% CI [0.950, 2.086]). As shown in Figure 11a, participants were more likely to reject negative EV gambles than accept positive EV gambles. Choices also differed significantly by RT quantile (*β* = 0.133, t(712) = 2.51, p = 0.01, 95% CI [0.029, 0.236]). There was no main effect of stimulus strength, but the interaction between stimulus strength and RT quantile was significant (*β* = 0.085, t(712) =2.57, p = 0.01, 95% CI [0.020, 0.149]). The interaction between gamble type and RT quantile was also significant (*β* =-0.103, t(712) = −3.13, p < 0.01, 95% CI [−0.167, −0.038]). None of the other interactions were significant.

**Figure 11.**
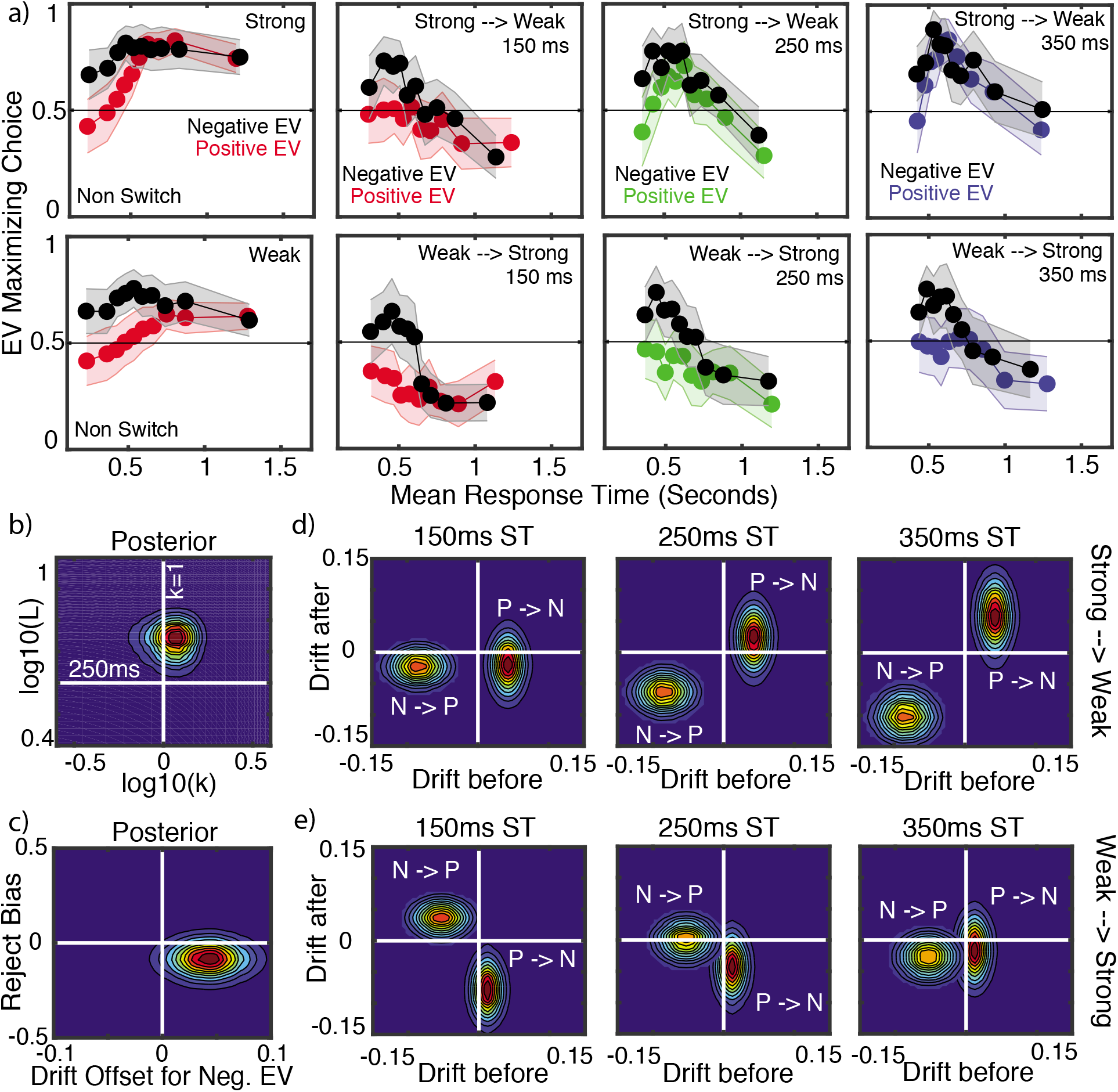
Experiment 4 Results. **Panel a)** The left most plots show the EV maximizing choice (accepting positive EV gambles and rejecting negative EV gambles) on non-switch trials for strong (top) and weak (bottom) information. The middle and right plots show the EV maximizing choices for the initial gamble (i.e., gamble presented at the start of the trial) in trials with responses after the switch time. Initially negative EV gambles are in black and initially positive EV gambles are in red (150 ms switch time), green (250 ms switch time), and blue (350 ms switch time). The strong to weak trials are on the top row and the weak to strong trials are on the bottom row. Choices are plotted for nine evenly space RT quantiles. Error band shows the 95% confidence interval. **Panel b)** Joint posterior distribution for the hyper means for the urgency ratio (*k*) and leakage (*L*) parameters. **Panel c)** Joint posterior distribution for the negative EV gamble drift offset and reject bias parameters. **Panes d, e)** Joint posterior distribution for the hyper means for the drift rates before and after the switch for the six switch conditions (three switch times along two different strength conditions). Plotting conventions are the same as in Figure 7 except that P and N refer to positive and negative EVs respectively.

For the analysis of the switch trials, we examined EV maximizing choices for the initial gamble (i.e., gamble presented at the start of the trial) in trials with responses after the switch time. For example, if the initial gamble is positive (i.e., more orange than blue), then the EV maximizing choice is ‘accept’ before the switch and ‘reject’ after the switch (in this case the grid changes to have more blue than orange and the new EV is negative). Thus, if participants are quickly adapting to the changed information, we should see a decrease in accept (reject) decisions when moving from a positive to negative (negative to positive) EV gamble. This is similar to the analyses in Experiments 1-3 where we examined the choices for the first majority color in trials with responses after the switch time (in this case, we should see a decrease in choices for the first majority color after the switch). We fit 16 different GLMMs, examining the inclusion of various fixed effects (i.e., a null model, models with only a single factor, models with two factors, models with three factors, and a model with all four factors). All models included an intercept. The model with all four factors (gamble type, stimulus strength, switch time, and RT quantile) had the lowest BIC and AIC values. The best fitting model showed that choices differed significantly by gamble type (*β* = 2.959, t(2144) = 4.12, p < 0.01, 95% CI [1.551, 4.367]). As shown in Figure 11a, participants were more likely to reject initially negative (but currently positive) EV gambles than accept initially positive (but currently negative) EV gambles. Choices also differed significantly by RT quantile (*β* = 0.367, t(2144) = 2.09, p = 0.037, 95% CI [0.022, 0.713]). In general, EV maximizing choices for the initial gamble declined with longer RTs. There was a marginally significant effect of switch time (*β* = 7.476, t(2144) = 1.83, p = 0.068, 95% CI [−0.553, 15.506]) and difficulty (*β* = 1.271, t(2144) = 1.89, p = 0.060, 95% CI [−0.051, 2.594]). There was a significant interaction between gamble type and RT quantile (*β* = −0.365, t(2144) = −3.27, p < 0.01, 95% CI [−0.584, −0.146]). None of the other interactions were significant.

We also examined detection ability using the results from Block 21. One participant was removed from this analysis because his / her hit and false alarm rates were both equal to 1, meaning that this participant responded “change” on all trials. The mean hit rate was 0.55 and the mean false alarm rate was 0.49. As in the previous experiments, discriminability (average d’ = .16) was significantly above zero, implying that participants could detect changes in this task (t(25) = 2.27, p = 0.032).

## Modeling Results

The general structure of the model used to assess the properties of the underlying decision process for this experiment is similar to that in Experiment 3 with two important differences. We once again account for (and estimate) the potential presence of urgency and leakage as well as allow for differing drift rates before and after the change of information. However, we account for the value based nature of this task and the potential effects of risk attitude and loss aversion (Tversky & Kahneman, 1992). First, we note that it is difficult to distinguish between aversion to risk as opposed to aversion to loss in our task because participants made decisions about single gambles offering both a positive and negative payoff (being risk averse in our task means avoiding the possibility of loss). Thus, we will we not draw strong conclusions about the difference between these two mechanisms. That said, there are at least two different ways that loss aversion / risk aversion can influence cognitive processes in our task.

First, participants might have an initial bias to reject any gamble with a possible loss, even before the gamble is known. This type of bias (which we call a “Reject Bias”) is best captured by a response bias parameter (z) that shifts the start-point of evidence accumulation towards the reject threshold (reflecting a general propensity to avoid risks).

The second way loss aversion / risk aversion can manifest is by modulating the processing of information. This type of bias is best modeled through the drift rates for negative expected value gambles. Specifically, we incorporate a “Drift Offset” parameter *DO* that affects the drift rates on negative EV gambles as follows. Consider two fixed stimulus trials (no change during the trial) with the same proportionality (difficulty), one majority orange (i.e., positive EV) and the other majority blue (i.e., negative EV). In the absence of associated value, we would expect that drifts for these stimuli to be the same (*μ_orange_* = *μ* and *μ_blue_* = –*μ*). However, if loss or risk aversion affects the processing of information, then this may not be the case. In order to account for asymmetries between positive and negative EV gambles, we incorporate a drift rate offset for the negative EV stimulus according to (*μ_orange_* = *μ* and *μ_blue_* = –*μ* – *DO*).

Now, consider a trial that switches from strong to weak with basic drift rates *μ_s,b_* (strong, before) and *μ_w,a_* (weak, after). In a positive → negative EV scenario the drift rates would transition from *μ_s,b_* → –*μ_w,a_* – *DO*. In a negative → positive EV scenario the drift rates transition from *μ_s,b_* – *DO* → –*μ_w,a_*. Here we are explicitly defining the upper boundary of the DDM process to correspond to a “gamble” response. We note that the reject bias and drift offset parameters are assumed to be independent of the duration and strength of the stimuli. Thus the model here contains only two more parameters than that associated with Experiment 3. This model allows us to both address the same questions investigated in Experiments 1-3 while controlling for potential biases introduced by the value based nature of this task.

Results of fitting this model to data (Figure 11) show broadly similar results as before with a few important differences (quality of model fit shown in Figure 8). The urgency ratio again appears to be moderate (k=1.5). However, the leakage appears to be faster, with a timescale of 180 ms. This result indicates that people may respond to changes of information more quickly in this value based scenario (although more research is needed to fully examine this possibility). That said, the urgency and leakage are broadly in the same ranges as found in Experiments 1-3. We once again also find that early information attenuates the perception of late information in both a strength and duration independent way, consistent with Experiments 1-3.

The value based nature of this task introduces both a start point “Reject Bias” as well as “Drift Offset” (Figure 11c). Specifically, we find that the reject bias is less than zero (posterior mean = −0.007, which is 8% of the threshold difference), indicating a stimulus independent preference to reject the gamble. This is consistent with prior work showing an initial bias for rejection when applying an evidence accumulation model to risky decision-making (Zhao, Walasek, & Bhatia, 2018). Such an initial bias might reflect a global-level of loss (or risk) aversion developed in the task. We also found that the “Drift Offset” is greater than zero (posterior mean = 0.04, where the base drift rates are on the scale of 0.01 – 0.1 in size), which results in drift rates that favor rejection of negative EV gambles more than acceptance of positive EV gambles, in line with loss aversion.

## Conclusions

Experiment 4 extends the previous findings in perceptual decision-making to a value-based task involving decisions about risky gambles. Results showed similar estimates for urgency and leakage as in Experiments 1-3, although leakage was faster in this task. Importantly, this experiment shows that information processing is not only dependent on the strength and duration of early information, but also on its value. Negative gambles resulted in steeper drift rates than equivalent positive gambles. As a result, early negative information had a larger impact on later positive information than vice versa.

### General Discussion

The recent debate about the properties of the cognitive processes responsible for fast perceptual decisions has centered around two hypotheses: 1) that neural architectures accumulate or integrate evidence over time until a decision threshold is met and 2) that time varying urgency modulates the weighting of evidence over time with more weight placed on later evidence. The majority of the studies on this topic, which, in aggregate paint a mixed picture, have sought to determine which of these mechanisms is most likely based on observations. We believe that not only is this approach a “false dilemma” (aka an “either-or fallacy”), but that it also overlooks the complexities associated with evidence leakage, which influences the decision process in conjunction with integration and urgency. Further, we believe that when a changing information paradigm is used to test these issues, as is necessary because the testing of dynamic properties is very difficult with static stimuli, a further issue arises, temporal dependence, where early arriving information changes the effect of different later arriving information.

In order to remedy these issues, we proposed a generalization of the Urgency Gating Model (UGM; Cisek et al., 2009) that accounts for potentially different “Urgency Signals”, the presence of leakage, and temporal dependencies in information processing. The urgency signal, as it was originally proposed, is effectively an evidence weight that increases over time, and which implements a tradeoff between speed and accuracy in a decision. However, most analyses of the UGM to date have assumed a restricted form for this urgency signal (a zero intercept, or *b* = 0 in the nomenclature of this paper), which fundamentally restricts the interpretation of the UGM’s scope and properties. This restriction was relaxed in Thura et al. (2014) and Thura and Cisek (2016a); however, in those studies, leakage was removed from the model and decision thresholds were fixed rather than estimated. Our generalization relaxes this restriction without making others, allowing for a wider class of urgency signals where a single parameter—the “Urgency Ratio” *k*—quantifies the relative influence of urgency versus integration. In extreme limits of this parameter corresponding to either flat or highly sloped urgency signals, this generalized model takes the form of the standard integration (when a 0 leakage rate also holds) or standard urgency models, which have been the topic of this “false dilemma”. A focus on extremes neglects the spectrum of possibilities that fall between, and we believe this has led to apparently conflicting results in the current literature.

The generalized model, the gUGM, generates a two parameter spectrum of models that differ not just in urgency characteristics but also the level of evidence leakage, which has been neglected in many of the studies investigating the dynamical nature of decisions. Crucially, these are distinct and estimable elements of the decision system (when the gUGM is applied to a changing-information paradigm), which could potentially be modulated separately. This is particularly important to note since prior studies have suggested that urgency and leakage lead to different kinds of optimality. Urgency has been suggested (Drugowitsch et al., 2012; Malhotra et al., 2017a) to be an optimal policy when difficulty varies across decisions, though our analysis here suggests this optimality may be weak at best with a range of different urgency and threshold values leading to similar reward rates. Leakage on the other hand has been posited to be an important element of optimal decision policies when evidence varies within decisions (Radillo et al., 2017; Kilpatrick et al., 2018).

In order to assess people’s decision properties, we combined this generalized model with a changing evidence paradigm to perform parameter based inference. Parameter inference was used rather than model comparison for a few reasons. First, the model comparison approach forces a researcher to explicitly define the models to be compared. Those models could be, for example, the standard DDM and standard UGM models. However, as we have discussed, there is a two parameter model space here worth considering, spanned by the critical urgency ratio and leakage parameters. Taking a parameter inference based approach in this case allows us to determine which portion of that model space best accounts for data, rather than focusing only on the extremes of the model space.

We also note that this parameter inference based approach yields some benefits compared with existing, more qualitative methodologies that have been used in the past. A common past approach has been to generate predictions from competing hypotheses that can be used to distinguish between them, and then run experiments to test those predictions. For example, Cisek et al. (2009) utilized this approach in their original article discussing the UGM. This approach, however, requires that the predictions being used to compare theories are actually distinguishing, which can be difficult to prove in practice. As detailed in Appendix B, our results demonstrate that the data on which the original support for the UGM were based (Cisek et al., 2009) do not distinguish between integration and urgency hypotheses when evidence is defined in a conventional manner, leading to inconclusive results (see Figure 12). Similarly, Hawkins, Wagenmakers, et al. (2015) previously suggested that response time skew is diagnostic of urgency versus integration with positive skew indicating integration is dominant while lack of skew is associated with urgency. However, as detailed in Appendix C, revisiting this result using this gUGM framework demonstrates that urgency and integration dominated models can make overlapping predictions, and thus skew is not always an effective differentiator of the two theories. In particular, while strong positive skew cannot be produced by pure urgency gating, weaker skew can be produced by both theories (see Figure 13).

**Figure 12.**
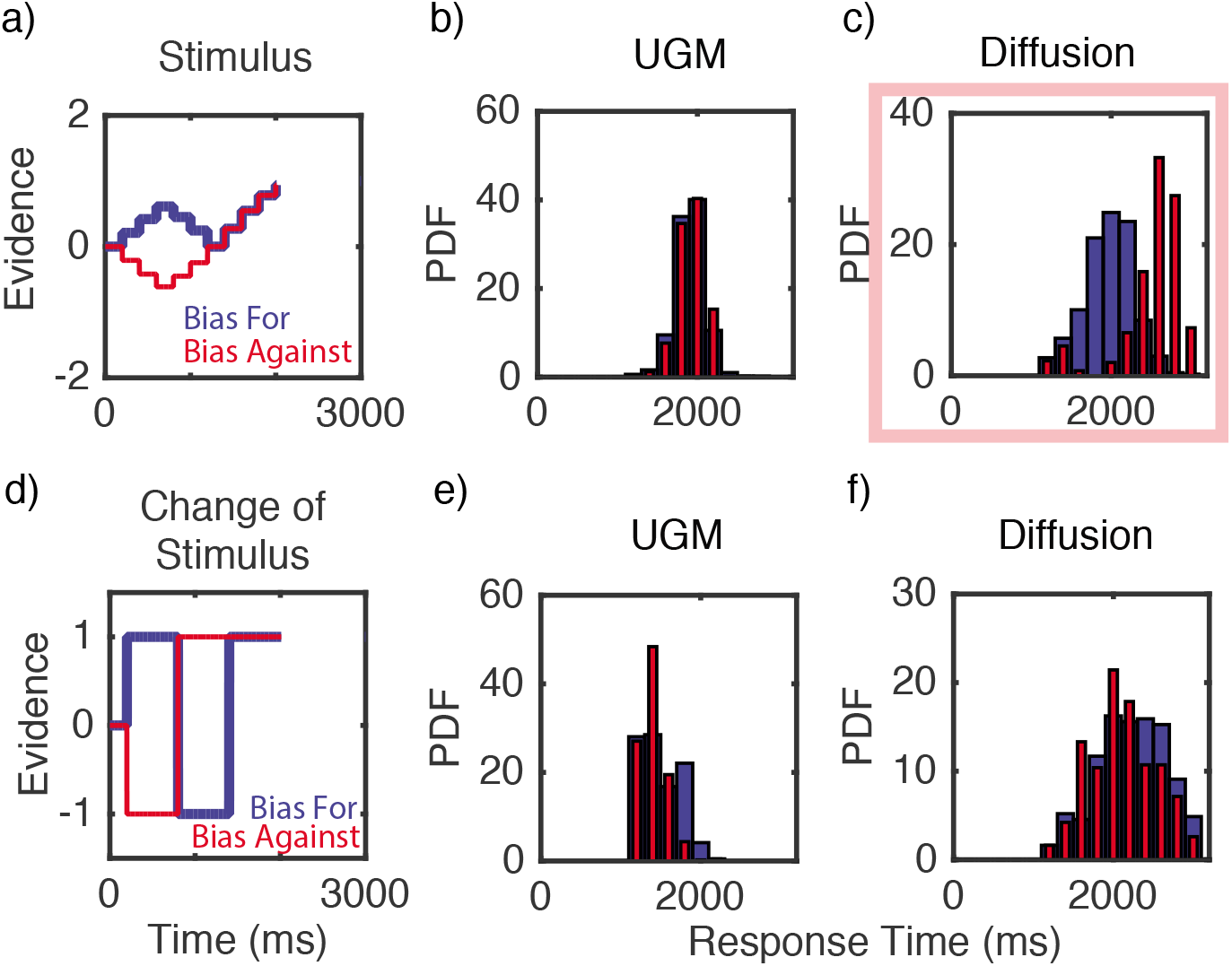
Influence of evidence interpretation on results. **Panels a, d)** Graphical depiction of two types of evidence signals for the “Bias for / Bias against” trial type considered in Cisek et al. (2009). In the terms of that publication, (a) would represent the success probability at a particular time, which depends on all information on the screen. (d) is effectively the derivative of that signal, depicting the change of stimulus. **Panels b, e)** Response time distributions simulated from the UGM with evidence signals (a, d) respectively. **Panels c, f)** Response time distributions simulated from the diffusion model with evidence signals (a, d) respectively. Panels (b, c) are exact reproductions of panels from Figure 7 in Cisek et al. (2009). The red square in (c) indicates qualitative disagreement with experimental results from Cisek et al. (2009).

**Figure 13.**
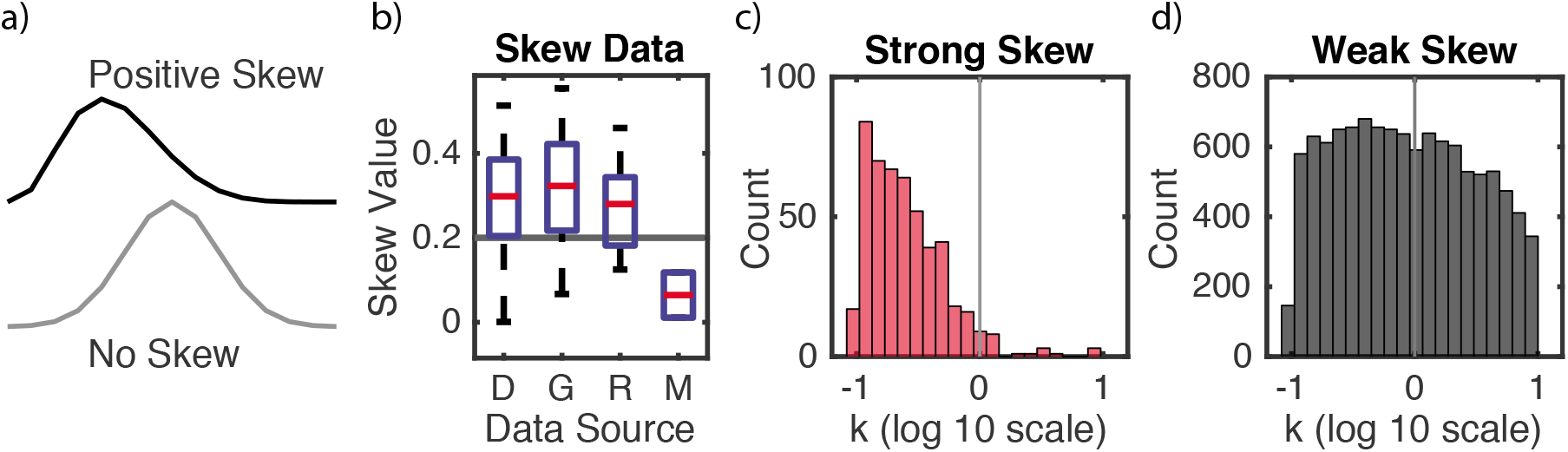
Influence of Urgency on Response Time Skew. **Panel a)** Schematic of RT distributions with positive and no skew. **Panel b)** Skew data calculated from RT distributions for 4 different data sets: D = human (Holmes et al., 2016), G = flashing grid data (Holmes & Trueblood, 2017), R = human (Ratcliff & McKoon, 2008), M = non-human primate (Roitman & Shadlen, 2002). Note that for data set M, there were only two non-human primate participants and thus the data has no error bars. The horizontal line is what we take to be the dividing line between weak (below) and strong (above) skew values. **Panels c, d)** To determine the correspondence between skew and urgency strength, 3000 parameter sets were generated in the parameter space of the generalized UGM using latin hyper cube sampling. Those with unreasonable RT distributions were discarded. For the remaining distributions, the skew value of the RT distribution was calculated. C and D respectively show the distribution of skew values that give rise to strong and weak skew respectively. For this study, the values of parameters were sampled from the interval *k, L, a* ∈ [0.1,10] with the remaining parameters fixed (*v* = 1, *σ* = 0.4).

The parameter inference based approach avoids these pitfalls, although it does of course have its own challenges. Foremost, the models being used for inference must be parametrically identifiable. Simulation studies here demonstrated that standard, static information paradigms where the information content in a stimulus is fixed during the course of a trial is insufficient to identify the parameters of the gUGM. However, when dynamic, changing information stimuli are used the model does become identifiable. In retrospect, this seems sensible. Time varying urgency and leakage are dynamic properties of the decision process. Urgency changes how information is weighted over time and leakage influences the persistence of information over time. This result thus suggests that dynamically changing stimuli are more effective at probing the dynamic properties of decisions than static stimuli. In light of these results we developed a changing information experimental paradigm to measure the dynamical properties (urgency ratio, leakage rate, and temporal drift rate dependencies) of the decision process.

Motivated by prior results in Holmes et al. (2016), we used our flashing grid paradigm to explicitly test how changes in duration, strength, and value of information influences decisions, and to assess how the decision process copes with those changes. We found three main results of broad interest. First, time varying urgency appears to be a part of the decision process but at a moderate level of *k* ~ 1. This is consistent with the results in Thura et al. (2014), where a broad range of 0.5-5 was found, though this study did not consider the potential effects of leakage or temporal dependence in information processing. Importantly, *k* ~ 1 is right at the dividing line of urgency and accumulation dominated scenarios, which may be one reason that prior studies focusing only on the two extreme ends of this spectrum have led to ambiguous and contradictory results.

Second, leakage appears to be present with a time constant of 200 – 250ms, which is significant, but on the slower (and hence weaker) end of the range that has typically been assumed in prior studies where leakage was fixed independent of the data (Table 1). These results were found to be consistent across three separate experiments involving purely perceptual decisions. The same moderate level of urgency, and slightly faster leakage, was also found in an experiment requiring value-based decisions. These results are also consistent with those of Holmes et al. (2016), who examined a paradigm in which the direction of random dot motion stimuli changed while decisions were being made. Their model did not allow for leakage, but it did allow for a delay in the onset of the effect of the change, which can mimic the effects of leakage, and they found a substantial delay of the order of several hundred milliseconds. Hence, it appears that leakage may be an important factor across a range of stimuli and decisions types.

Third, information processing in this changing information task is relative rather than veridical. When information changes, initial evidence influences the perception of later information in a duration, strength, and value dependent manner. In Experiments 1-3, we show that longer duration and stronger early information has a larger influence on subsequent drift rates. In Experiment 4, we extend this to show that the value of early information also influences later information, with early negative information having a greater impact on later positive information than vice versa. This temporal dependence is also consistent with the results reported by Holmes et al. (2016). Because that study used motion stimuli (i.e., a RDM task), they suggested the temporal dependence could be due to a motion after effect. The results we found here with non-motion stimuli however suggest that temporal dependence may be a more general feature of decisions where information changes.

## Conclusion

In sum, this paper presents the first (to our knowledge) comprehensive application of a modeling framework that can account for three key dynamic elements of the decision process: urgency, leakage, and temporal dependencies in information processing. As we illustrate, all three mechanisms are important and necessary in characterizing decisions about changing information. Across four experiments in both perceptual and value-based decision-making domains, we show that urgency and leakage are present and that the duration, strength, and value of early information alters the processing of later information. In the future, we hope that researchers interested in questions about the dynamics of information processing in decision-making will consider using a comprehensive modeling framework, such as the one presented here, because simplifying approaches that omit one or more of the key mechanisms we discussed can lead to false conclusions.

## Appendix A Modeling and parameter estimation methods

Following Turner et al. (2013), we fit all models as hierarchal models in a Bayesian framework using a Differential Evolution Markov Chain Monte Carlo (DE-MCMC) procedure. The purpose of using this procedure rather than a more typical MCMC is to account for the correlation between parameters that is common in accumulator type models (Turner et al., 2013) that lead to high rejection rates and poor efficiency. We briefly describe the implementation details specific to this context and refer the reader to Ter Braak (2006), Turner et al. (2013), and Storn and Price (1997) for a more complete discussion.

The model we fit to this data set is a hierarchal extension of the gUGM model described previously. Given the simple structure of this experiment where all participants come from a single population, we assume that the hierarchal structure consists of a single group. Based on this, we make the standard assumption that individuals are members of a normally distributed population and assign a normal prior for each individual level parameter

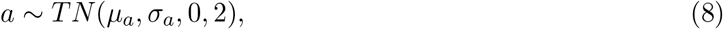

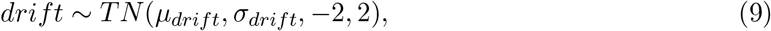

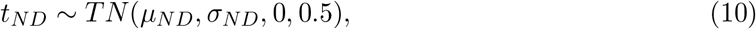

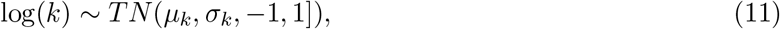

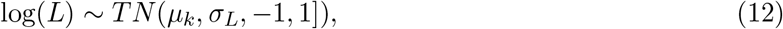

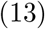

where *TN*(*μ, σ, a, b*) refers to the normal distribution with mean *μ* and standard deviation *σ* truncated to the interval [*a, b*]. All drift rate parameters were subject to the same priors. Additionally, log normal priors were used for leakage and urgency ratios since, for example, values of the urgency ratio of 0.1 and 10 represent symmetric extremes that are well captured in the logarithmic domain. We further specify the following mildly informative priors for the hyper mean

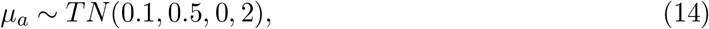

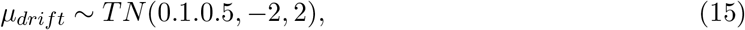

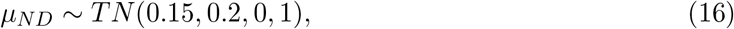

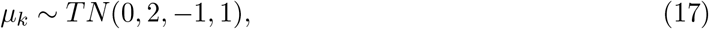

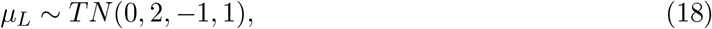

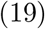

and hyper standard deviation parameters respectively

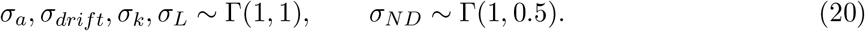

Here, Γ(*a, b*) refers to the gamma distribution with shape and rate parameters *a* and *b* respectively.

In the case of experiment 4 where an additional start point bias was included, the following priors and hyper-priors were used

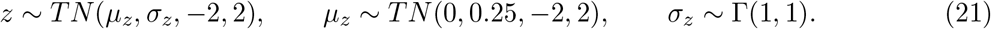

Similarly, the drift offset parameter was given the following priors

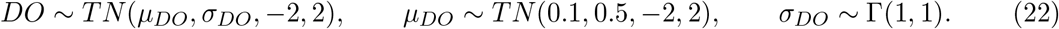

To fit this hierarchal model, we utilize the PDA method (Holmes, 2015; Turner & Sederberg, 2014b) embedded into the DEMCMC framework with 10,000 simulated samples used to approximate the likelihood function for each condition and a time step of *dt* = 5ms. To assess quality of fit, we plot the actual and predicted choice probability and mean RT for all participants (see Figure 8).

## Appendix B Revisiting the Motivation for Time Varying Urgency

Here we revisit the original supporting evidence for the UGM hypothesis (Cisek et al., 2009). One prominent methodology for attempting to discern whether decision processes are more urgency or integration dominant has been to determine qualitatively distinct predictions of the two hypotheses and design experimental paradigms test for that difference. In particular, one of the earliest works attempting to assess the presence of these factors is Cisek et al. (2009), which claimed to reveal inconsistencies between EAM predictions and behavioral results with time varying stimuli. While this approach is in principle sound, it is necessary to demonstrate that the predictions on which experiments are based actually distinguish between hypotheses. In this section, we revisit the results from Cisek et al. (2009) to demonstrate the pitfalls of this approach.

The original motivation for the development of the UGM was the suggestion that EAM’s such as the diffusion decision model (DDM) produce choice / RT predictions that are inconsistent with observations when stimulus information changes over time (Cisek et al., 2009; Thura et al., 2012). To support this conclusion, the authors utilized a behavioral task where a central group of dots contained within a circle on the screen discretely move to initially empty circles on the left or right, one by one, at fixed intervals. In this task, the participants goal was to predict which circle (left or right) would eventually end up with the most dots (see Cisek et al., 2009 for further detail). For present purposes the two critical binary choice experimental conditions that were used to distinguish between EAM and UGM hypotheses were “Bias for” (BF) and “Bias against” (BA) trials. In these trials initial sensory evidence (i.e., the first few objects to leave the center circle) briefly favors the choice that will (BF), or will not (BA), later be consistently favored. Their critical observation was that cumulative success rates were clearly different, but RTs were indistinguishable for these two trial types. They key claim that we focus on here is that EAMs produce RT distributions inconsistent with data, whereas the UGM is consistent with data.

Our re-analysis (Figure 12) indicates this conclusion depends on a non-standard interpretation of what constitutes “evidence”. Their assumption was that sampled evidence was associated with “success probability”, which is calculated based on the complete knowledge of the full state of the screen and how many dots were left, right, and center (their Eqn. 30). A more common assumption, made to our knowledge in all earlier work on decisions on the direction of motion (Ratcliff & McKoon, 2008; Roitman & Shadlen, 2002), is that evidence sampled at a given time is associated with object motion (e.g., that a dot moves to the left or to the right, see Figure 12d for the evidence signal considered). In the words of Thura et al. (2012), “a good decision policy should only integrate sensory signals to the extent that they provide novel information”. Cisek et al. (2009) did consider a form of novel evidence, however the implementation took an unusual form where a delta function like shot of evidence was present only at the instant when the stimulus changed. Specifically, they assumed evidence was present for the 1ms after the stimulus changed followed by a 199ms window when evidence is assumed to be 0. We consider a minor adjustment of this model that instead assumes the evidence signal takes the form shown in Figure 12d, which is a smooth signal consisting of consistent evidence for one alternative followed by consistent evidence for the other. We re-analyzed the UGM and EAM hypotheses with this type of evidence (Figures 12 (e, f)) and performed the same qualitative comparison of the data that they performed. Results show that both UGM and EAM produce qualitatively indistinguishable RT distributions for BF / BA trials, both consistent with data. We also qualitatively compared choice probabilities on these trial types choice / RT predictions on their other trial types (easy / hard) and found both models to again be consistent with data. These results demonstrate the value of using quantitative fitting to test models rather than relying solely on qualitative predictions, which can be difficult to concretely validate as distinguishing.

## Appendix C Is Response Time Skew Diagnostic of Decision Strategy?

In Hawkins, Wagenmakers, et al. (2015) it was suggested that the statistical skew of RT distributions provides a diagnostic tool for determining whether urgency or integration is responsible for decisions. Specifically, that positive skew (i.e., a preponderance of long relative to short RTs) is associated with integration, and no skew (i.e., symmetry, see Figure 13(a)) is associated with urgency. We are, however, unaware of a systematic study that assesses the robustness of this association.

As the generalized UGM allows for the presence of both mechanisms at different levels of strength, we assessed the association between the aforementioned urgency ratio, which measures the relative importance of urgency / integration, and skew. Figures 13 (c,d) provides an analysis of the correlation between cognitive strategies (integration / urgency) and response time skew. For this analysis, 100,000 unique parameter sets (*k, L, a*) were generated using a latin hyper cube design in a logarithmically spaced parameter domain (for (*k, L*) so that large (> 1) and small (< 1) values of these parameters are equally represented). For simplicity, the non-decision time (*t*_0_) was fixed at 300ms while the within trial stochastic noise magnitude was fixed at *s* = 0.1. For each parameter set, 1000 independent stochastic simulations were performed with a 1ms time step. The resulting data was weakly censored so that parameter sets generating unreasonable RT distributions with a standard deviation less than 100ms were removed from further analysis. This left 11,650 remaining parameter sets for which the value of the distributional skew was calculated. To set a reasonable dividing line between what represents a “skewed” versus “unskewed” RT distribution, we assessed the value of RT skew from four published perceptual decision making data sets, three from random dot motion tasks, two with human participants (R) Ratcliff and McKoon (2008), and (D) Holmes et al. (2016), an one with non-human primates (M) Roitman and Shadlen (2002), and a numerosity-judgement task (G) Holmes and Trueblood (2017). Based on this distribution of skew values (Figure 13b), we set a threshold of 0.2 to represent the dividing line (shown as the gray horizontal line) between strongly skewed and weakly skewed. For each of the 11,650 tested parameter sets, the simulation results were categorized as either a strongly or weakly skewed generating parameter set based on this demarcation line.

Figures 13 (c, d) show the frequency distribution of values of the urgency ratio associated with strong and weak skew respectively. They demonstrate that a strong positively skewed distribution is almost exclusively associated with a model where integration dominates urgency (i.e., *k* < 1). However, weak or no skew is not diagnostic, as both urgency and integration dominated models produce distributions with weak skew. Figures 13 (b) shows skew data from three human and one non-human primate studies. All of the human studies are consistent with integration dominance, with the non-human primate study being consistent, although not exclusively so, with urgency gating. One possible source of this difference between the skew of human and non-human response time skew is the context of the experiment. Whereas human behavioral experiments typically involve little training and relatively weak incentives, non-human experiments typically require a great deal of training and clear incentives. This could lead to differences in decision traits, for example, accuracy maximization versus reward rate maximization and interrogation versus free response paradigms. More generally, these results demonstrate how the gUGM can be used to systematically probe for associations between simple non-parametric empirical measures (skew in this case) and the relative importance of integration / urgency.

## Footnotes

1 Note that more participants were excluded in this experiment than Experiment 1. This is because the practice trials used for exclusion in Experiment 1 (i.e., block 2) were easier than the practice blocks in Experiment 2.

## Notes

WRH, JST, and NE were supported by NSF grant SES-1556325. AH was supported by ARC grant DP160101891. All authors contributed in a significant way to the manuscript and all authors approved the final manuscript.

